# Single-cell transcriptome analysis reveals CD34 as a novel marker of human sinoatrial node pacemaker cardiomyocytes

**DOI:** 10.1101/2024.09.06.611657

**Authors:** Amos A. Lim, Delaram Pouyabahar, Mishal Ashraf, Kate Huang, Michelle Lohbihler, Matthew L. Chang, Brandon M. Murareanu, Thinh Tran, Amine Mazine, Gary Bader, Zachary Laksman, Stephanie Protze

## Abstract

The sinoatrial node (SAN) regulates the heart rate throughout life. Failure of this primary pacemaker results in life-threatening, slow heart rhythm. Despite its important function, the cellular and molecular composition of the human SAN is not completely resolved. Particularly, no cell surface marker to identify and isolate SAN pacemaker cells has been reported to date. Here we used single-nuclei/cell RNA sequencing of fetal and human pluripotent stem cell (hPSC)- derived SAN cells and show that the SAN consists of three subtypes of pacemaker cells, including Core SAN, SAN, and Transitional Cells. Our study identified a host of novel Core SAN markers including MYH11, BMP4, and the cell surface antigen CD34. We demonstrate that sorting for CD34^+^ cells from cardiac hPSC differentiations enriches for SAN cells with a functional pacemaker phenotype. This novel SAN pacemaker cell surface marker is highly valuable for future hPSC-based disease modelling, drug discovery, cell replacement therapies, as well as the delivery of therapeutics to SAN cells *in vivo* using antibody-drug conjugates.

## Introduction

The sinoatrial node (SAN) is the primary pacemaker of the heart located at the boundary of the right atrium and the superior vena cava. The pacemaker cells of the SAN generate electrical impulses that initiate each of the three billion heart beats that a human heart experiences in a lifetime. SAN diseases can lead to a life-threatening decrease in heart rate, a condition known as bradyarrhythmia. The current standard treatment for SAN dysfunction is implantation of an electronic pacemaker device which, while lifesaving, has multiple downsides^1^. To improve treatment options, a better understanding of the SAN pacemaker cells and their molecular identifiers are needed.

The SAN is a small structure that is comprised of about 10,000 pacemaker cells and has therefore been hard to access and study. Additionally, it is a heterogenous tissue that contains multiple cell types, including fibroblasts, smooth muscle cells, endothelial cells, and neurons from the innervating autonomous nervous system, as well as different subtypes of pacemaker cardiomyocytes. In the human heart, these include the Core SAN pacemaker cardiomyocytes that express the well-established pacemaker genes *TBX3, TBX18, ISL1, HCN4*, and lack the expression of the pan-cardiomyocyte marker *NKX2-5*^2–4^. These core pacemaker cells are responsible for initiating the heartbeat. In addition, a transitional cell type, also referred to as paranodal cells, has been described that expresses both pacemaker and atrial cardiomyocyte genes^5–7^. These cells are thought to facilitate the conduction of the electrical signals from the SAN tissue into the atrial myocardium.

Bulk RNA sequencing has been performed to gain a better understanding of the molecular identity of the SAN pacemaker cardiomyocytes, but tissue resident non-cardiomyocytes resulted in the detection of off-target genes that are expressed in neurons or fibroblasts^4, 8, 9^. Single-cell-based sequencing technologies have revolutionized our ability to study tissue compositions at the transcriptome level. Single-cell RNA sequencing (scRNA-seq) of the mouse SAN uncovered several new SAN pacemaker cardiomyocyte-specific marker genes^9–11^. Most recently, Kanemaru & Cranley et al. dissected SAN tissue from adult hearts to generate the first single-cell transcriptome atlas of the human SAN, but no single-cell-based dataset of the developing human SAN has been established yet^12^.

Studies of the human SAN are hampered by the limited access to healthy primary tissue. To overcome this, investigators have turned to the human pluripotent stem cell (hPSC) system as an alternative source of SAN pacemaker cells^13^. Accessing these cells from patient-specific, induced pluripotent stem cells (iPSCs) enables modelling of SAN diseases and to gain deeper insights into disease mechanisms. Beyond that, hPSC-derived SAN cardiomyocytes are a source of functional pacemaker cells for the development of biological pacemakers that could overcome the downsides of electronic pacing. Several groups, including our own, have developed protocols for the directed differentiation of hPSCs into SAN-like pacemaker cells (SANLPCs)^14–18^. To isolate enriched populations of SANLPCs from the differentiation cultures, transgenic reporters have been employed, including NKX2-5:GFP for the negative selection of NKX2-5^-^ SANLPCs or SHOX2:GFP for the positive selection of SHOX2^+^ SANLPCs^14, 15, 18^. Identification of a cell surface antigen that is specifically expressed on SANLPCs would overcome the need to generate reporter lines and enable a straightforward identification and isolation of SANLPCs from differentiation cultures of any hPSC line. To date, a SANLPC specific cell surface marker has not been identified. To gain a better understanding of the cellular and molecular composition of the human SAN and identify potential SAN specific cell surface markers, we performed single-nuclei/single-cell RNA sequencing (sn/scRNA-seq) of human fetal SAN tissue and hPSC-derived SANLPCs. This allowed us, to show that hPSC-derived SANLPCs closely resemble fetal SAN pacemaker cells on the transcriptome level. Our analysis identified three subtypes of SAN pacemaker cells, including Core SAN, SAN, and Transitional Cells. Combining the fetal and hPSC-derived datasets, we established a shared list of Core SAN pacemaker cell specific genes. This list contains a host of novel SAN markers, most importantly the surface antigen CD34. We show that CD34 is specifically expressed by the pacemaker cardiomyocytes in human SAN tissue and by hPSC-derived SANLPCs, but not by other cardiomyocyte subtypes. Finally, we demonstrate that sorting for CD34^+^ cells from hPSC differentiation cultures enriches for SANLPCs with a functional pacemaker phenotype.

## Results

### Single-cell transcriptomic analysis of the human fetal sinoatrial node

To assess the molecular heterogeneity of the cells in the developing human SAN, we dissected the SAN tissue of a fetal heart (gestation week 19) and performed snRNA-seq (Fig. 1a). Unsupervised clustering identified 19 cell clusters representing cell types that are expected to be present in the heart, including cardiomyocytes as well as non-cardiomyocytes such as fibroblasts, smooth muscle, endothelial, endocardial and epicardial cells, neurons, and macrophages^9, 19^ (Fig. 1b, Supplementary Table 1). To obtain a better resolution of the cardiomyocyte subtypes, we performed subcluster analysis on the cardiomyocyte cell clusters. This resulted in eight cell clusters that expressed high levels of *TNNT2,* indicative of cardiomyocytes (Fig. 1 b, c). To identify the cardiomyocytes representing SAN pacemaker cells, we analyzed the expression of established SAN pacemaker genes (*TBX3, TBX18, SHOX2, ISL1, HCN4*)^2, 4, 5^. We detected one cell cluster expressing high levels of all these genes that we annotated as Core SAN cells. We identified a second cell cluster expressing these SAN genes but with lower expression levels of *TBX3* and *ISL1* that we annotated as SAN cells. Of note, these Core SAN and SAN cells did not express the cardiac transcription factor *NKX2-5*, which is a well-established characteristic of SAN pacemaker cells^2, 3^. In addition, three cell clusters representing Atrial cells were identified based on the expression of *NKX2-5*, the atrial marker *NPPA,* and *SCN5A*^4, 19^. As expected, we also identified a cluster of cells expressing both atrial and pacemaker genes, representing the myocytes of the Transition Zone between the pacemaker and atrial tissues^6, 7, 9^. Finally, two clusters of proliferating *MKI67* expressing cells were detected. Differential gene expression analysis showed that the Transition Zone cells were marked by the expression of *LRRC4C* (netrin-G1 ligand), *MYO16* (myosin 16), and *SLC24A3* (sodium/potassium/calcium exchanger 3), while SAN cells were marked by the expression of the long non-coding RNA *LINC01880*, *PDEA1* (phosphodiesterase 1A), and *CDH13* (cadherin 13) (Fig. 1d, Supplementary Table 2). We next focused our analysis on the Core SAN cells to identify novel markers of this population. The Core SAN cells expressed the highest levels of the established pan-pacemaker marker *TBX3*^2, 20^. In addition, the expression of other well-known pacemaker genes (*TBX18*, *SHOX2*, *ISL1*, *RGS6*, *VSNL1*, *CACNA1D*, *HCN1*, *HCN4*) was significantly enriched in this cell cluster, confirming a pacemaker phenotype^4, 8, 9, 21^. Gene ontology (GO) analysis for biological processes of the genes enriched in the Core SAN cells resulted in terms including regulation of heart rate, cardiac pacemaker cell differentiation, sinoatrial node development, and regulation of SAN cell action potential, further supporting the SAN pacemaker phenotype of these cells (Fig. 1e). Importantly, the top ten differentially expressed genes (DEGs) within the Core SAN cluster contained several novel genes not previously reported to be involved in SAN pacemaker cell function or development, including *TENM2* (teneurin transmembrane protein 2)*, PPFIA2* (protein tyrosine phosphatase receptor type F polypeptide interacting protein alpha 2)*, CNTN4/5* (contactin 4/5) *PRKG1* (protein kinase cGMP-dependent 1)*, SLIT2* (slit guidance ligand 2)*, DGKB* (diacylglycerol kinase beta), and *KCNIP4* (potassium voltage-gated channel interacting protein 4) (Fig. 1 d, f). Furthermore, the guanylate kinase-associated protein *DLGAP1*, recently described as a SAN specific gene in the mouse, was also contained in our top ten gene list^10^.

**Figure 1:**
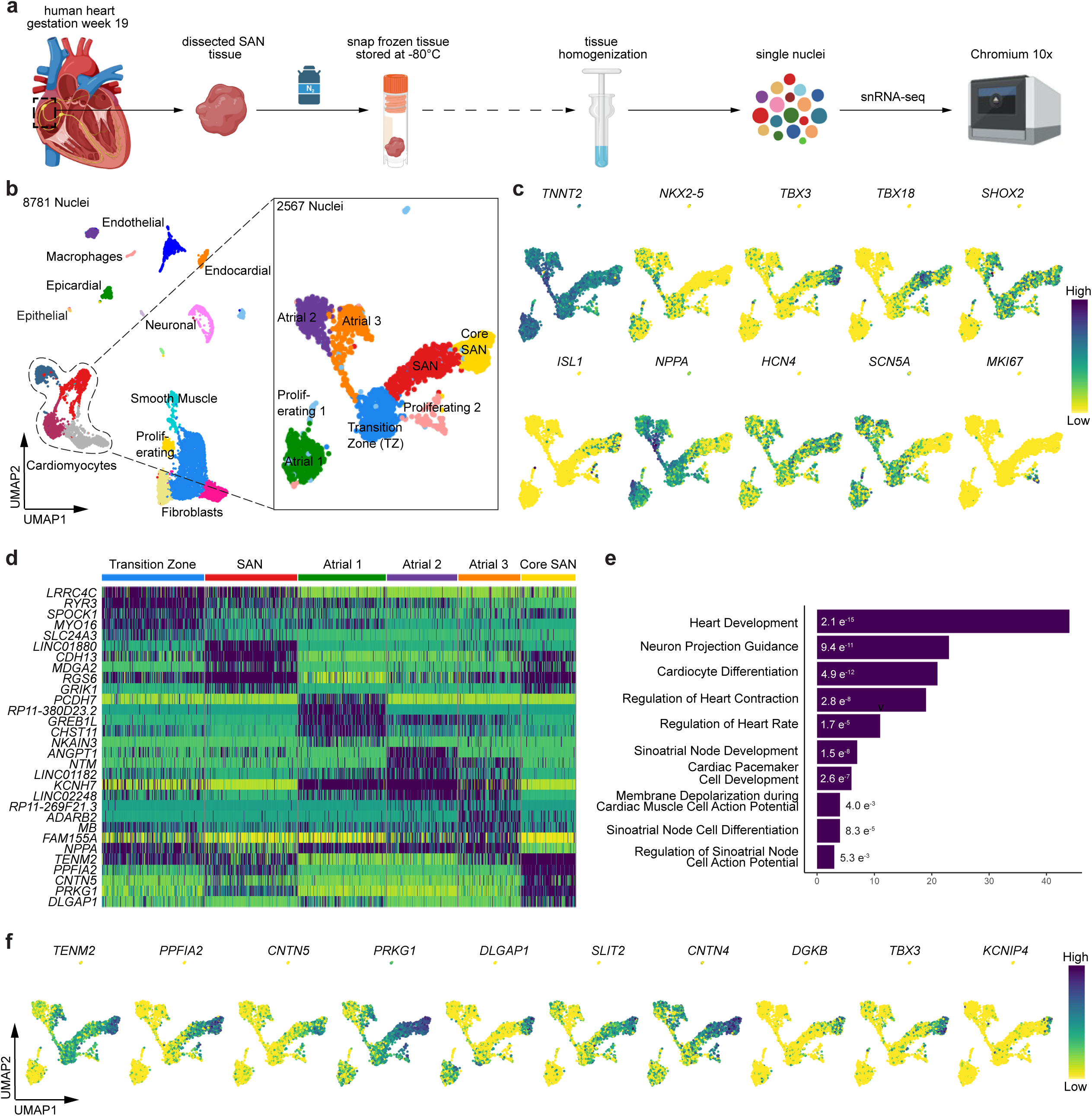
Single-nuclei RNA sequencing of fetal SAN tissue identifies markers of Core SAN pacemaker cells. **a** Schematic overview of SAN tissue dissection and processing for snRNA-seq on the 10x Genomics Chromium platform. **b** Uniform manifold approximation and projection (UMAP) of gestation week 19 fetal heart SAN tissue sample showing 19 cell clusters (left). Subclustering of the *TNNT2*^+^ cardiomyocytes showing 8 sub-clusters (right). **c** UMAPs of the subclustered cardiomyocytes showing the expression of the indicated genes. **d** Heatmap of the top 5 differentially expressed genes (DEGs) within the indicated cardiomyocyte subclusters. **e** Gene Ontology (GO) analysis (biological processes) of all genes enriched in the Core SAN cluster. **f** UMAPs showing the top 10 DEGs of the core SAN cluster.

### hPSC-derived SAN like pacemaker cells transcriptionally closely resemble fetal SAN pacemaker cells

We next wanted to compare the expression profiles of the fetal SAN cells to hPSC-derived SANLPCs generated using our previously reported protocol^14^ (Fig. 2a). This protocol generates a mixed population, containing both NKX2-5^-^ SANLPCs (50%) and NKX2-5^+^ cardiomyocytes (30%), that we used for scRNA-seq analysis. As shown in Fig. 2b, 11 cell clusters were identified, including fibroblasts, epithelial, and epicardial cells, as well as cardiomyocytes. To specifically compare the expression between the cardiomyocyte subtypes, we next subclustered the *TNNT2*^+^ cardiomyocytes. Analysis of the expression of established SAN pacemaker markers *TBX3, TBX18, SHOX2, ISL1,* and *HCN4*^2, 4, 5^ identified a Core SAN cluster that was negative for *NKX2-5* expression (Fig. 2c). Like in the fetal SAN tissue, a second *NKX2-5*^-^ cardiomyocyte cluster was identified that expressed all the pacemaker markers, but lower levels of *TBX3*, that we annotated as SAN cells. Analysis of expression of the atrial marker *NPPA* together with working cardiomyocyte markers *NKX2-5* and *SCN5A* identified a cluster containing Atrial cardiomyocytes. The hPSC-cultures also contained Transition Zone cells that expressed both pacemaker and atrial genes. In addition, two clusters of proliferating *MKI67* expressing cardiomyocytes were identified. To further confirm the identity of the hPSC-derived cardiomyocyte clusters, we scored the cells in the hPSC-derived dataset with the DEG lists of the fetal tissue dataset^22^ (Fig. 2d). Applying the top 200 DEGs from the fetal Core SAN cluster clearly identified the Core SAN cluster in the hPSC dataset. Using the same approach, the identities of the SAN, Transition Zone and Atrial cell clusters were confirmed. These findings indicate that the NKX2-5^+^ cells within the hPSC-derived population represent Atrial and Transition Zone cells while the NKX2-5^-^ cells represent Core SAN and SAN pacemaker cells.

**Figure 2:**
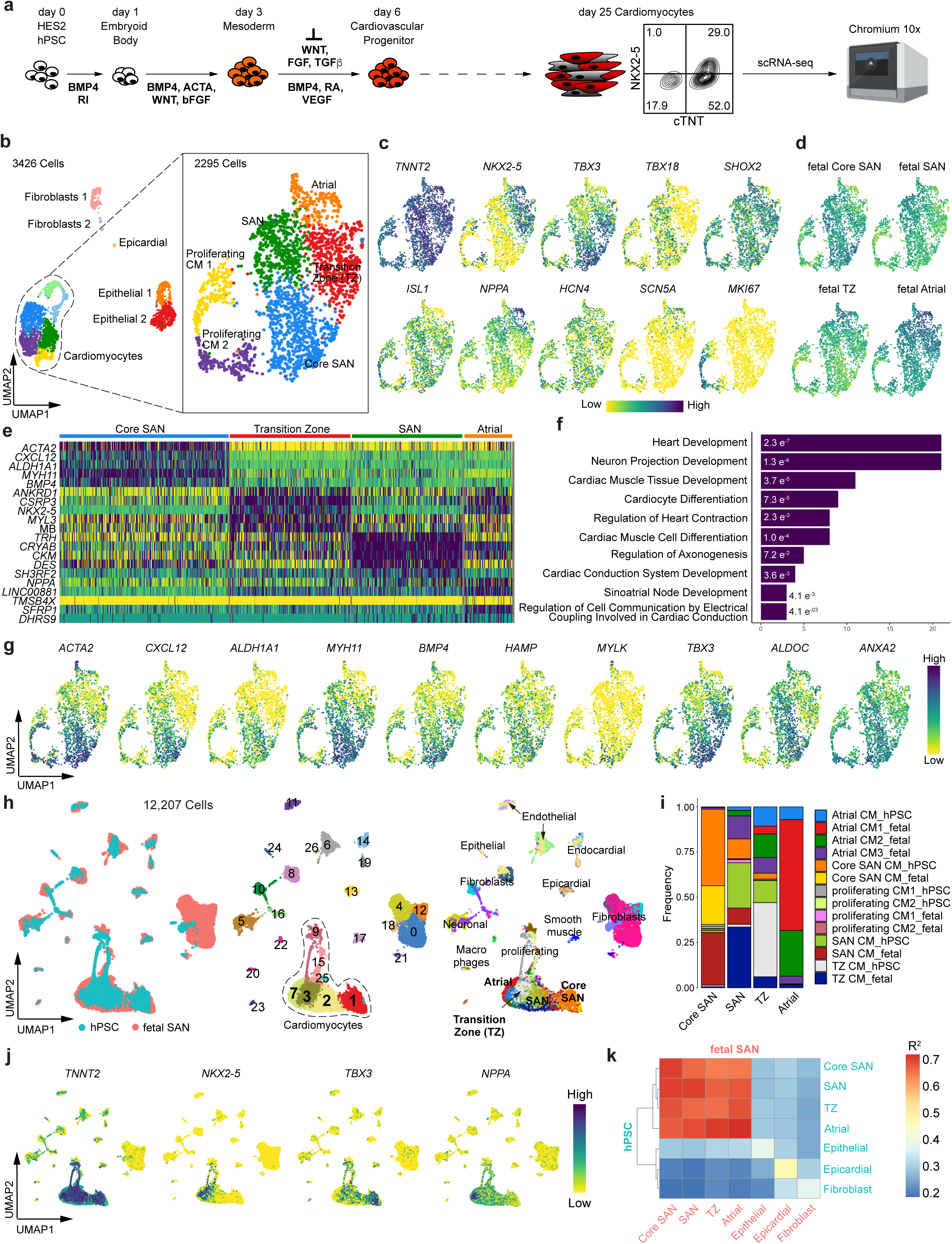
Single-cell RNA sequencing of hPSC-derived SANLPCs reveals transcriptomic similarities to fetal SAN pacemaker cells. **a** Schematic overview of SANLPC differentiation protocol and sample processing for scRNA-seq on the 10x Genomics Chromium platform. **b** UMAP of day 25 HES2-derived SANLPCs showing 11 cell clusters (left). Subclustering of the *TNNT2*^+^ cardiomyocytes showing 6 sub-clusters (right). **c** UMAPs of the subclustered cardiomyocytes showing the expression of the indicated genes. **d** UMAPs showing signature score distribution for the DEGs of the indicated fetal SAN cell types. **e** Heatmap of the top 5 DEGs within the indicated cardiomyocyte subclusters. **f** GO analysis (biological processes) of all genes enriched in the Core SAN cluster. **g** UMAPs showing the top 10 DEGs of the core SAN cluster. **h** UMAPs showing Harmony integration of the fetal SAN snRNA-seq data and the hPSC scRNA-seq data labelled by source dataset (left), cluster number (centre), and assigned cell types (right). **i** Stacked bar graph showing the frequency of each cell type from both fetal and hPSC datasets in the indicated cell clusters. **j** UMAPs of integrated fetal SAN and hPSC datasets showing the expression of the indicated genes. **k** Spearman correlation between selected clusters from the fetal SAN and hPSC datasets. p < 0.05 for all correlations.

Differential gene expression analysis identified expression patterns that distinguish the different hPSC-derived cardiomyocyte subtypes. Transition Zone cells were marked by the expression of *ANKRD1* (ankyrin repeat domain 1, cardiac muscle), *CSRP3* (cysteine and glycine rich protein 3, cardiac LIM protein), and *MYL3* (myosin light chain 3), while SAN cells were marked by the expression of *TRH* (thyrotropin releasing hormone), *CKM* (creatine kinase, muscle) and *DES* (desmin) (Fig. 2e, Supplementary Table 3). The Core SAN cells were enriched in the expression of previously established conduction system and SAN pacemaker markers (*TBX3, VSNL1, LBH, CPNE5*)^4, 8, 9, 20, 23^. GO term analysis identified an enrichment of genes involved in regulation of heart contraction, cardiac conduction system development, and sinoatrial node development, further supporting the SAN pacemaker phenotype of the cells contained in the Core SAN cluster (Fig. 2f). Interestingly, the top ten DEGs of the Core SAN cells contained genes that have not previously been reported to be specifically expressed in SAN pacemaker myocytes including *ACTA2* (actin alpha 2, smooth muscle), *ALDH1A1* (aldehyde dehydrogenase 1a1), *HAMP* (hepcidin antimicrobial peptide), *MYLK* (myosin light chain kinase, smooth muscle), and *ALDOC* (aldolase c, brain-type) (Fig. 2g). The list also contained the C-X-C motif chemokine ligand *CXCL12*, the smooth muscle myosin heavy chain isoform *MYH11*, the BMP signaling ligand *BMP4,* and the annexin family member *ANXA2* which have been previously detected in transcriptomic analysis of mouse and human SAN cells^4, 8, 12, 21^.

To further compare the hPSC-derived cells and the fetal SAN cells, we used Harmony integration and combined the two datasets (Fig. 2h-j, Supplementary Fig. 1a, b)^24^. This analysis showed that the hPSC-derived cardiomyocytes clustered together with the fetal cardiomyocytes. Within the integrated cardiomyocytes we identified clusters of Core SAN, SAN, Transition Zone and Atrial cardiomyocytes as described in the individual datasets. Importantly, the Core SAN cluster contained both hPSC-derived and fetal tissue-derived Core SAN cells (Fig. 2i, Supplementary Fig. 1a). The vast majority of the Core SAN cells from the hPSC and fetal datasets (74% and 94%) clustered together in this merged Core SAN cluster. The SAN, Transition Zone, and Atrial clusters similarly contained both hPSC-derived and fetal-derived cells of the respective cardiomyocyte subtype. However, along with the matching subtypes a more heterogenous contribution of other cardiomyocyte subtypes was detected in these clusters. Spearman correlation analysis further demonstrated that the hPSC-derived Core SAN cells are most similar to the cells of the fetal Core SAN (Fig. 2k). Taken together, this molecular analysis indicates that the hPSC-derived Core SAN pacemaker cells closely resemble the Core SAN pacemaker cells found in the developing human heart.

### Identification of shared Core SAN pacemaker markers

To identify a list of marker genes that can be used to identify SAN pacemaker cells *in vitro* and *in vivo*, we next compared the DEGs of the Core SAN cells from the hPSC and fetal datasets. This analysis identified 36 genes that were significantly enriched in both populations (Fig. 3a-d, Supplementary Fig. 2a-c). Signature scoring analysis showed that these 36 genes clearly identify the Core SAN cell clusters in the hPSC and fetal data. The gene list contains the established cardiac conduction system and SAN markers *TBX3, TBX5, VSNL1, LBH, CPNE* ^2, 4, 8, 9, 20, 23^ as well as *GNAO,* a subunit of the G-protein signal- transducing complex, that has been recently described to be expressed in hPSC-derived SAN-like cells^21^. In addition, this shared list contains a number of novel genes that have not been previously associated with human SAN function or development including; (i) contractile apparatus and cytoskeletal genes: *MYH11* (myosin heavy chain 11), *ACTB* (actin beta), *WIPF3* (WAS/WASL interacting protein family member 3); (ii) genes involved in BMP signaling: *BMP4* (bone morphogenetic protein 4), *BMP5* (bone morphogenetic protein 5), *ID3* (inhibitor of DNA binding 3); (iii) genes encoding for calcium binding proteins involved in cellular signalling: *ANXA2* (annexin 2), *NECAB1* (N-terminal EF-hand calcium binding protein 1), *SPOCK1* (proteoglycan 1); (iv) genes encoding for regulators of transmembrane calcium ion-currents: *RRAD* (Ras related glycolysis inhibitor and calcium channel regulator), *PRKG1* (protein kinase cGMP dependent 1); (v) and genes classically known to be expressed in neurons and glia cells that play a role in axon growth and guidance: *BASP1* (brain abundant membrane attached signal protein 1), *PPFIA2* (protein tyrosine phosphatase receptor type F polypeptide interacting protein alpha 2), *EFR3B* (EFR3 homolog B), *SLIT2* (slit guidance ligand 2), *LYPD6* (LY6/PLAUR domain containing 6). Importantly, the list also contains five genes encoding membrane spanning proteins that could serve as SAN pacemaker cell surface markers that have not been previously associated with SAN cells: *CD34* (CD34 antigen), *CADM1* (cell adhesion molecule 1), *EFNB2* (ephrin B2), *NTRK2* (neurotrophic receptor tyrosine kinase 2), and *ELAPOR2* (endosome-lysosome associated apoptosis and autophagy regulator family member 2).

**Figure 3:**
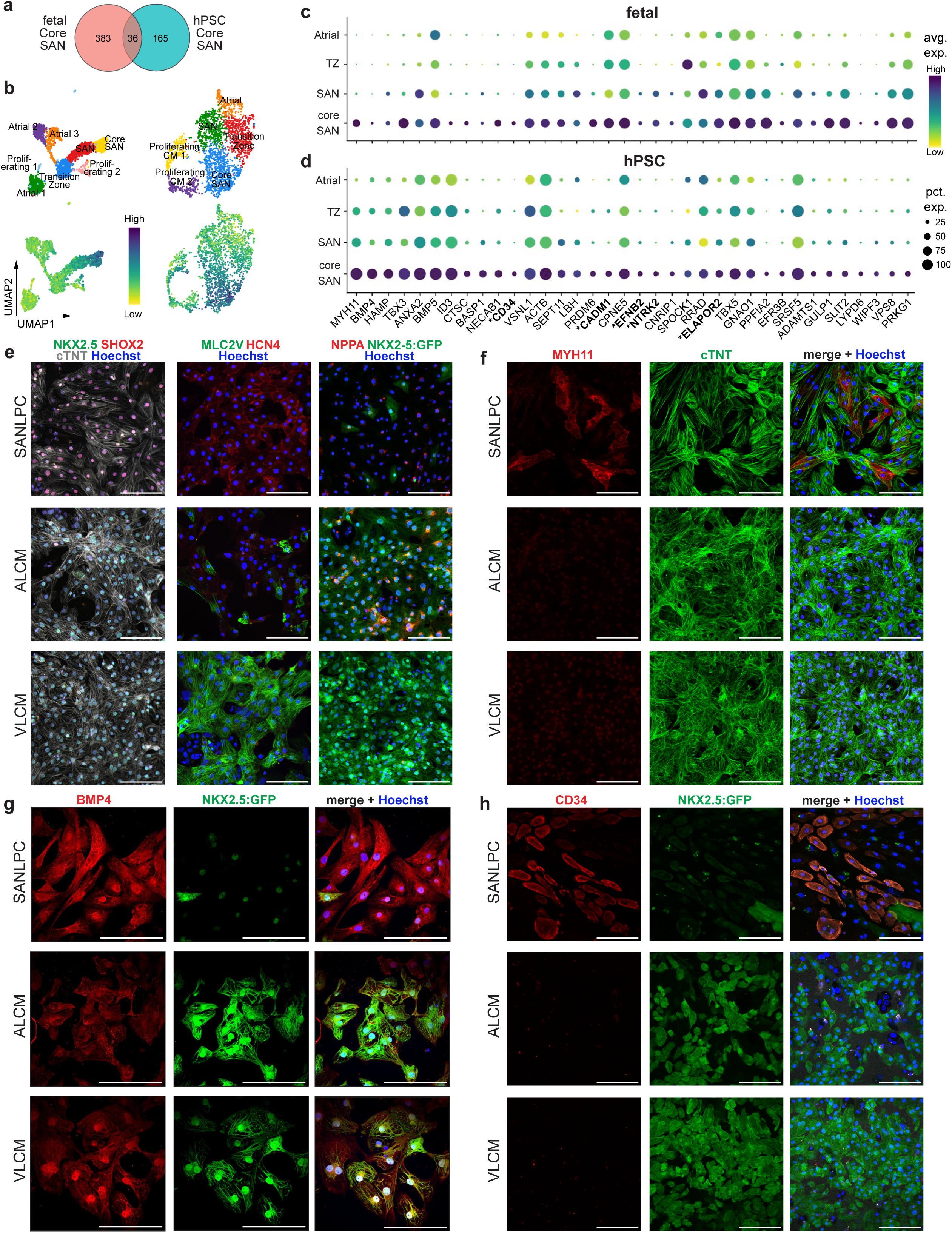
Comparison of fetal and hPSC-derived SAN cells identifies shared Core SAN markers. **a** Venn diagram of the Core SAN markers identified in the fetal and hPSC datasets. **b** UMAPs of *TNNT2*^+^ cardiomyocytes showing the assigned cell types (top row) and the signature score distribution (bottom row) of the shared 36 Core SAN marker genes in the fetal (left) and hPSC datasets (right). **c, d** Dotplots showing the expression of the conserved core SAN markers in the fetal (**c**) and hPSC (**d**) datasets. **\***indicates genes encoding for membrane spanning proteins. **e-h** Immunofluorescent staining of day 25 hPSC-derived SANLPCs, atrial-like cardiomyocytes (ALCMs), and ventricular-like cardiomyocytes (VLCMs) for NKX2-5 and SAN pacemaker transcription factor SHOX2, ventricular contractile apparatus protein MLC2V and pacemaker ion channel HCN4, NKX2-5:GFP and atrial protein NPPA (**e**), Core SAN marker MYH11 (**f)**, Core SAN marker BMP4 (**g**), and Core SAN marker CD34 (**h**). Cells were counterstained with cTNT to identify cardiomyocytes and Hoechst to visualize all cells. Scale bars, 100μm.

To determine if these markers are conserved between species, we compared our human Core SAN marker list to the previously published scRNA-seq data of the mouse SAN from Goodyer et al. ^9, 25^ (Supplementary Fig. 3a-c). As expected, the established SAN markers *TBX3, TBX5, VSNL1, LBH,* and *CPNE5* were conserved between human and mouse. However, only half of the human Core SAN genes from our list were specifically expressed in mouse SAN pacemaker cells. The other half was higher expressed in atrial cardiomyocytes of the mouse or expressed at very low levels. These observations prompted us to assess whether the SAN specific genes identified in the mouse SAN scRNA-seq dataset are conserved in the human SAN^9^. Out of the 50 genes that we analyzed, the majority (56%) were either not detected in the human Core SAN cells or expressed at higher levels in human atrial cardiomyocytes. Notably, the list of genes also contained mouse SAN pacemaker cell surface markers^25^, and only 13 out of 25 were conserved in the human SAN. Taken together, this analysis shows that, while key markers are conserved between species, there are numerous markers that are specific to the human SAN. This highlights the importance of analyzing human cells and tissues.

### MYH11, BMP4, and CD34 are specifically expressed in human SAN pacemaker cardiomyocytes

We selected three Core SAN genes identified in our study to further validate them as SAN markers. We chose the top two differentially expressed genes *MYH11* and *BMP4*, which have previously been detected in transcriptomic studies of SAN pacemaker cells but have not been further validated on the protein level^4, 8, 21^. In addition, we chose the surface antigen CD34 due to its promising potential application as a SAN cell surface marker. We first focused on hPSC-derived cardiomyocytes and used previously reported protocols to generate SANLPCs, atrial-like cardiomyocytes (ALCMs) and ventricular-like cardiomyocytes (VLCMs) ^14, 26^. RT-qPCR analysis and immunofluorescent staining confirmed the phenotype of the different cardiomyocyte subtypes. As expected, SANLPCs expressed SHOX2 and HCN4, ALCMs expressed NKX2-5 and NPPA, and VLCMs expressed NKX2-5 and MLC2V (Fig. 3e, Supplementary Fig. 4 a, b). Importantly, immunostaining for MHY11, BMP4 and CD34 showed that these proteins are specifically expressed by SANLPCs but not ALCMs and VLCMs (Fig. 3f-h).

To validate the expression of these new SAN markers in primary tissue, we dissected the SAN of fetal hearts (gestation week 17-20) and prepared longitudinal sections for immunostaining (Fig. 4a, Supplementary Fig. 5a)^27^. We used the expression of TBX3, SHOX2 and HCN4 to identify the fetal SAN pacemaker cells, expression of NPPA to identify adjacent atrial cardiomyocytes, and expression of cTNT to distinguish cardiomyocytes from non-cardiomyocytes (Fig. 4aI, II). Immunostaining for MYH11 clearly labelled a subset of HCN4^+^cTNT^+^ SAN pacemaker cardiomyocytes but did not label any atrial cardiomyocytes (Fig. 4aIII). As expected, MYH11 was also detected in the smooth muscle cells of blood vessels contained in the sections. BMP4 expression also colocalized with HCN4^+^cTNT^+^ SAN pacemaker cells, where it was higher than in the adjacent atrial tissue (Fig. 4aIV). CD34 specifically labelled cTNT^+^ SAN cardiomyocytes but not atrial cardiomyocytes (Fig. 4aV). As expected, we detected CD34^+^NKX2-5^-^ SAN cardiomyocytes representing the Core SAN pacemaker cells identified in our transcriptional analysis (white arrowheads). In addition, we detected a few CD34^+^NKX2-5^+^ (green arrowheads) and CD34^-^NKX2- 5^-^ (yellow arrowheads) pacemaker myocytes in the SAN. Of note, CD34 is also expressed by endothelial cells, which most likely explains the detection of some CD34^+^ cells in the atrial tissue. Co-staining with CD31 and cTNT confirmed that these CD34^+^ cells are indeed CD31^+^ endothelial cells and that only SAN cardiomyocytes but not atrial cardiomyocytes stain positive for CD34 (Supplementary Fig. 5b).

**Figure 4:**
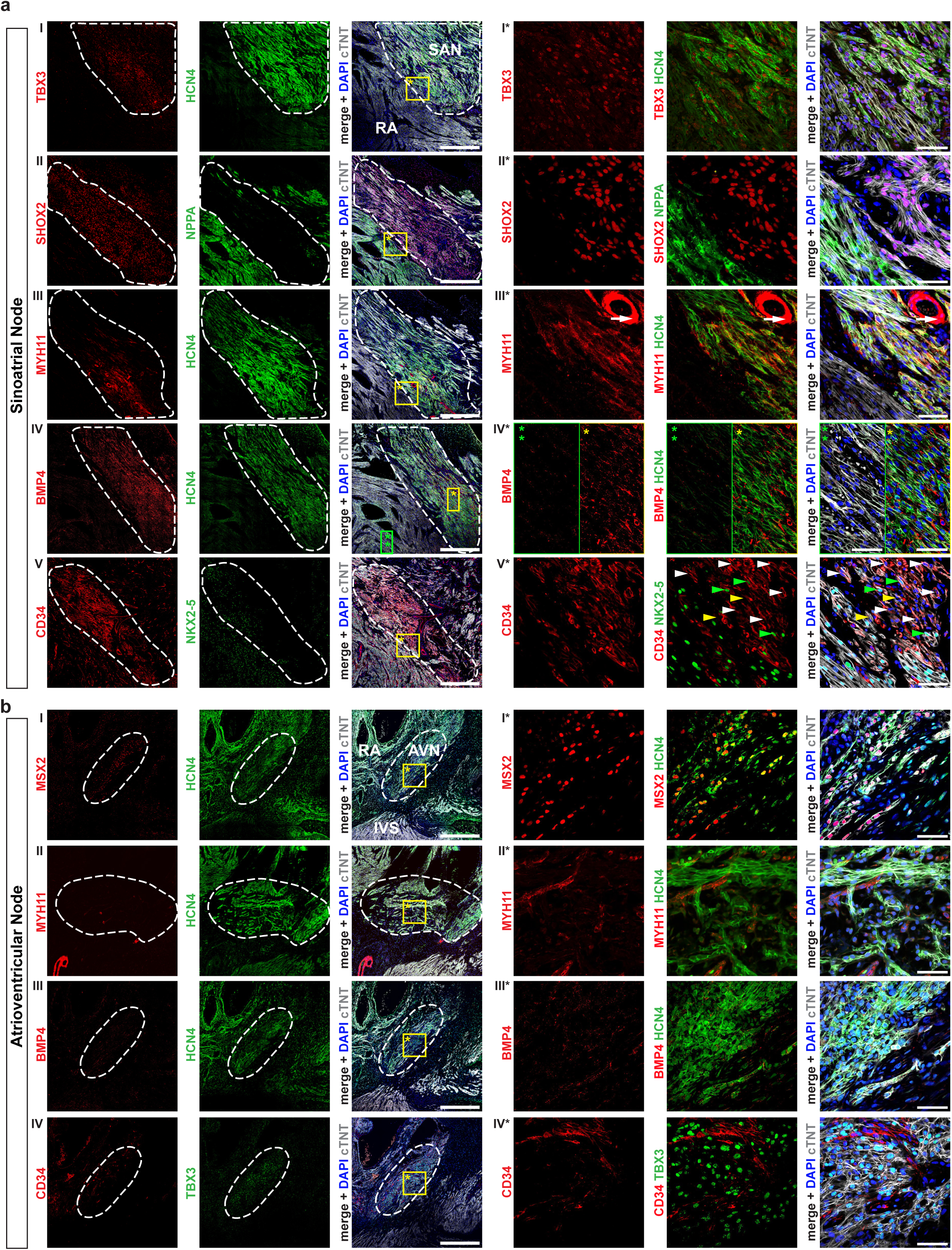
MYH11, BMP4 and CD34 are specifically expressed in human SAN pacemaker cardiomyocytes. **a** Immunofluorescent staining of gestation week 19 fetal human SAN for: pacemaker transcription factor TBX3 and pacemaker ion channel HCN4 (I), SAN pacemaker transcription factor SHOX2 and atrial protein NPPA (II), core SAN marker MYH11 and HCN4 (III), Core SAN marker BMP4 and HCN4 (IV), and Core SAN marker CD34 and NKX2-5 (V). Images represent consecutive sections of SAN tissue. White dashed line outlines the SAN. Yellow and green boxes indicate location of high magnification insets shown on the right marked with *. White arrows in III* indicate MYH11^+^ smooth muscle cells of a blood vessel. Arrowheads in V* indicate the following cell types: white, CD34^+^NKX2-5^-^; green, CD34^+^NKX2-5^+^; yellow, CD34^-^NKX2- 5^-^. **b** Immunofluorescent staining of gestation week 17 fetal human AVN for: AVN transcription factor MSX2 and HCN4 (I), Core SAN marker MYH11 and HCN4 (II), Core SAN marker BMP4 and HCN4 (III), and Core SAN marker CD34 and TBX3 (IV). Images represent consecutive sections of AVN tissue. White dashed line outlines the AVN. Yellow box indicates location of high magnification insets shown on the right marked with *. Tissue sections were counterstained with cTNT to identify cardiomyocytes and DAPI to visualize all cells. Scale bars, 500μm (left) and 50μm in the high magnification insets (right). IVS, interventricular septum; RA, right atrium

Our computational analysis suggested that CD34 is specific to human SAN pacemaker cells, because its expression was not detected in mouse SAN pacemaker cells (Supplementary Fig. 3b)^9^. To further validate this, we performed immunostaining for CD34 on sections of mouse SAN tissue (postnatal days 0-3). These experiments confirmed that SHOX2^+^TBX3^+^HCN4^+^ mouse SAN cardiomyocytes do not express CD34 (Supplementary Fig. 6a, b).

To determine if expression of MYH11, BMP4 and CD34 is specific to the hearts primary pacemaker, the SAN, or whether these proteins are also expressed by the secondary pacemaker, the atrioventricular node (AVN), we analyzed sections of the atrioventricular junctional region of fetal hearts (gestation week 17-20) (Supplementary Fig. 5a). AVN pacemaker cardiomyocytes were identified based on the expression of the AVN markers TBX3, MSX2, and the lack of expression of the SAN marker SHOX2 (Fig. 4bI, IV, Supplementary Fig. 5cI, II). NPPA and MLC2V expression identified adjacent atrial and ventricular tissues, respectively. Neither MYH11 nor BMP4 were expressed in AVN cardiomyocytes (Fig. 4bII, III). Similarly, no CD34^+^cTNT^+^ were found in AVN tissue (Fig. 4bIV, Supplementary Fig. 5cIII). The CD34^+^cTNT^-^ cells that were detected likely represent a CD34^+^ mesenchymal/fibroblast cell population that has previously been reported in multiple studies^19, 28, 29^. Taken together, these findings show that MYH11, BMP4, and CD34 are specifically expressed by the primary pacemaker cardiomyocytes of the SAN, but not by atrial, ventricular, nor AVN cardiomyocytes.

### Isolation of hPSC-derived SAN-like pacemaker cells based on CD34 expression

To determine if CD34 could be used for the isolation of SANLPCs from hPSC differentiation cultures, we first carried out flow cytometric analysis of CD34 expression. The HES3 NKX2-5^egfp/w^ reporter line was used for these experiments which allowed us to identify NKX2-5:GFP^-^ cardiomyocytes as SANLPCs and NKX2-5^+^ cardiomyocytes as VLCMs or ALCMs^14, 30^. Analysis within the fraction of SIRPA^+^CD90^-^ cardiomyocytes^31^ at day 25 of differentiation showed that only SANLPCs are marked by CD34 while VLCMs and ALCMs do not express CD34 (Fig. 5a, b, Supplementary Fig. 7a). Within the SANLPC differentiation cultures, the majority of the NKX2-5^-^ cells expressed CD34 (54 ± 4%). Notably, a small fraction of NKX2-5^+^ cells contained in the SANLPC cultures also expressed CD34 (17 ± 2%), an observation consistent with our findings in the fetal heart tissue sections. To identify the earliest timepoint of CD34 expression in SANLPC progenitors, we performed a time course analysis (Fig. 5c, d). The first SIPRA^+^CD90^-^ cardiomyocytes detected at day 8 of differentiation did not express CD34. By day 10, a small population of NKX2-5^-^ cardiomyocytes started to express CD34 and by day 16, most of the NKX2-5^-^ cardiomyocytes expressed CD34. The proportion of CD34^+^NKX2-5^-^ cardiomyocytes continued to increase until day 50 at which point 81 ± 4% of NKX2-5^-^ cardiomyocytes expressed CD34 (Supplementary Fig. 7b, c). The first NKX2-5^+^CD34^+^ cardiomyocytes were detected at day 16. The proportion of NKX2-5^+^CD34^+^ cardiomyocytes also increased over time, but expression levels of CD34 remained lower than in the NKX2-5^-^ cardiomyocytes at all timepoints analyzed. Accordingly, analysis at day 50 showed that 64 ± 6% of NKX2-5^-^ cells and 20 ± 3% of NKX2-5^+^ cells expressed high levels of CD34 (CD34^++^), similar to the proportions of CD34^+^ cells observed at day 25. Importantly, day 50 VLCM and ALCM populations remained CD34 negative, confirming that CD34 is exclusively expressed on SAN pacemaker cardiomyocytes (Supplementary Fig. 7d, e).

**Figure 5:**
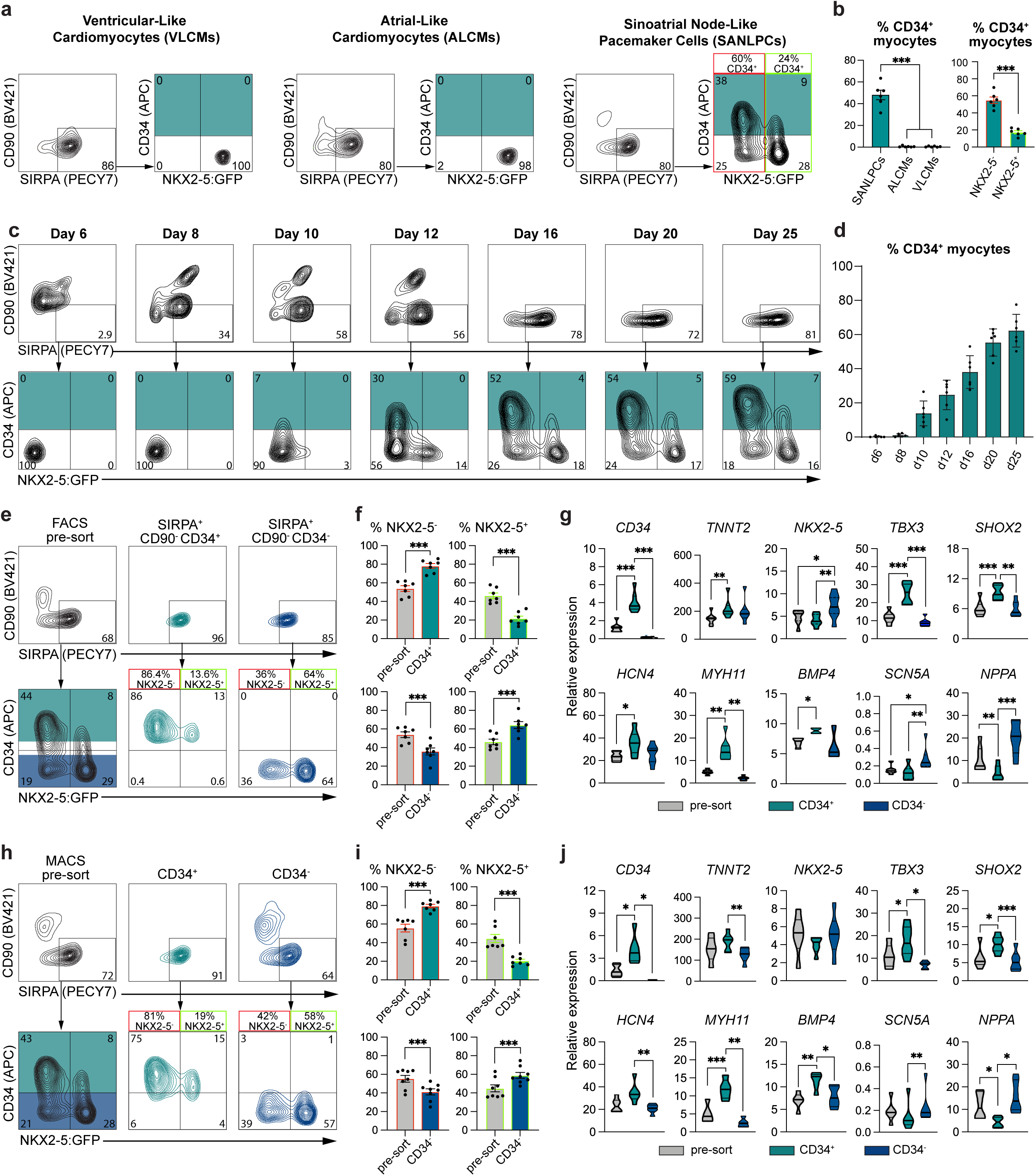
FACS and MACS sorting for CD34^+^ cells enriches for SANLPCs. **a** Flow cytometric analyses at day 25 of CD34 and NKX2-5:GFP expression in SIRPA^+^CD90^-^ cardiomyocytes in VLCMs, ALCMs, and SANLPCs. **b** Bar graphs summarizing the expression of CD34 in myocytes as shown in (a) in the indicated differentiation cultures (left) and within the NKX2-5^-^ and NKX2-5^+^ fraction of SANLPC differentiation cultures (right) (n = 6). **c** Flow cytometric analyses of CD34 and NKX2-5:GFP expression in SIRPA^+^CD90^-^ cardiomyocytes at indicated time points throughout the differentiation. **d** Bar graph summarizing the expression of CD34 in myocytes as shown in (c) (n = 6). **e** Flow cytometric analyses of CD34 and NKX2-5:GFP expression in SIRPA^+^CD90^-^ cardiomyocytes before and after FACS at day 25. Teal shading indicates CD34^+^ and blue shading indicates CD34^-^ FACS sorting gates. **f** Bar graphs summarizing the proportion of NKX2-5^-^ and NKX2-5^+^ cells in presort, CD34^+^, and CD34^-^ FACS sorted samples (n = 7). **g** RT-qPCR analysis of the expression of the indicated genes in presort, CD34^+^, and CD34^-^ FACS sorted samples (n = 7). Values represent expression relative to the housekeeping gene TBP. **h** Flow cytometric analyses of CD34 and NKX2-5:GFP expression in SIRPA^+^CD90^-^ cardiomyocytes before and after MACS at day 25. Teal shading indicates CD34^+^ and blue shading indicates CD34^-^ MACS sorting gates. **i** Bar graphs summarizing the proportion of NKX2-5^-^ and NKX2-5^+^ cells in presort, CD34^+^, and CD34^-^ MACS sorted samples (n = 8). **j** RT-qPCR analysis of the expression of the indicated genes in presort, CD34^+^, and CD34^-^ MACS sorted samples (n = 5-6). Values represent expression relative to the housekeeping gene TBP. Statistical analysis was performed using two-sided unpaired t-test when comparing two samples (b, f, i) and one-way ANOVA followed by Bonferroni’s post hoc test when comparing >2 samples (b, g, j): *p < 0.05, **p < 0.01, ***p < 0.001 vs indicated sample. Error bars represent SEM.

To test if it is possible to enrich for NKX2-5^-^ SANLPCs based on CD34 expression, we isolated SIRPA^+^CD90^-^CD34^+^ and SIRPA^+^CD90^-^CD34^-^ cardiomyocytes by fluorescence-activated cell sorting (FACS) and analyzed the populations for NKX2-5:GFP expression (Fig. 5e, f). We perform these experiments at day 25 of differentiation because at this timepoint the majority of NKX2-5^-^ cells expressed CD34 while the majority of NKX2-5^+^ cells were still CD34 negative. As shown in Figure 5e and 5f, the CD34^+^ sorted population was significantly enriched for NKX2-5^-^ cells and significantly depleted of NKX2-5^+^ cells compared to pre-sort. The opposite trend was observed for the CD34^-^ sorted population. RT-qPCR, analysis showed significantly enriched expression of SAN genes (*TBX3, SHOX2, HCN4, MYH11, BMP4*) and decrease expression of atrial/working cardiomyocyte genes (*SCN5A, NPPA*) in the CD34^+^ sorted cells compared to pre-sort and CD34^-^ sorted cells (Fig. 5g). To determine if the CD34^+^NKX2-5^-^ phenotypeis stable over time, we cultured the cells for an additional 25 days. Flow cytometric analyses at days 1, 10 and 25 post-sort revealed that the cells maintained CD34 expression and did not upregulate *NKX2-5* expression (Supplementary Fig. 7f, g).

For disease modelling and cell therapy applications, large cell numbers of SAN pacemaker cells will be required, thus prompting us to test whether magnetic-activated cell sorting (MACS) for CD34^+^ cells can also enrich for SANLPCs. Interestingly, MACS for CD34^+^ cells enriched for SIRPA^+^CD90^-^ cardiomyocytes without the need for additional antibodies (Fig. 5h-j, Supplementary Fig. 7h). Notably, the CD34^+^ sorted cells were also significantly enriched in NKX2-5^-^ cardiomyocytes and depleted of NKX2-5^+^ cells compared to pre-sort. Accordingly, RT-qPCR analysis showed increased expression of SAN pacemaker genes in the MACS sorted CD34^+^ cells and decreased expression of working cardiomyocyte genes compared to pre-sort and CD34^-^ sorted cells (Fig. 5j). The CD34^+^ population contained a low frequency (< 5%) of CD31^+^ endothelial cells that are known to contaminate hPSC-derived cardiomyocyte differentiations (Supplementary Fig. 7i, j). These cells were no longer detectable following 5 days of culture, likely because our culture conditions did not support their survival.

To confirm that this CD34-based isolation of NX2-5^-^ SANLPCs can be applied to other hPSC lines, we repeated the experiments using the HES2 hESC line (Supplementary Fig. 8a-f). First, we confirmed that CD34 is specifically expressed by HES2-derived SANLPCs but not by ALCMs and VLCMs. Next, we showed that MACS sorting for CD34^+^ cells results in a significant enrichment of NKX2-5^-^ cardiomyocytes and depletion of NKX2-5^+^ cardiomyocytes. In addition, CD34^+^ sorted cells showed higher expression of SAN pacemaker genes and reduced expression of working cardiomyocyte genes, thus confirming the enrichment for SANLPCs. Taken together, these experiments demonstrate that CD34 can be universally used to identify NKX2-5^-^ SANLPCs and isolate them from cardiac hPSC-differentiation cultures.

### CD34^+^NKX2-5^+^ and CD34^-^NKX2-5^-^ cardiomyocytes transcriptionally represent SAN pacemaker cells

In addition to the CD34^+^NKX2-5^-^ SAN pacemaker cells we detected some CD34^+^NKX2-5^+^ and CD34^-^NKX2-5^-^ cardiomyocytes in the fetal SAN (Fig. 4aV) and hPSC-derived SANLPC cultures (Fig. 5a, b). Furthermore, sorting for CD34^+^ cells also isolated a small proportion of CD34^+^NKX2-5^+^ cells (Fig. 5e-i). To characterize the identity of both of these cardiomyocyte populations in more detail, we performed Cellular Indexing of Transcriptomes and Epitopes by sequencing (CITE-seq) on day 25 of SANLPC differentiation using a barcoded CD34 antibody (Supplementary Fig. 9a, b)^32^. This enabled us to read out protein level CD34 expression in parallel to the whole transcriptome of the cells. Unsupervised clustering identified 11 cell clusters with the majority representing cardiomyocytes. Small clusters of fibroblasts, epithelial cells, and epicardial cells were also detected with a similar distribution as in our original hPSC dataset (Fig. 6a). Subclustering of *TNNT2^+^* cardiomyocytes identified the same cardiomyocyte subtypes as before, including *NKX2-5*^-^*TBX3*^high^ Core SAN cells, *NKX2-5*^-^*TBX3*^mid^ SAN cells, *NKX2-5*^+^*NPPA*^+^ Atrial cells, and Transition Zone cells that express both pacemaker and atrial genes (Fig. 6a, b, Supplementary Table 4). Signature scoring analysis using the top 200 DEGs of the fetal SAN dataset as a reference confirmed the identity of these cardiomyocyte subclusters (Fig. 6c).

**Figure 6:**
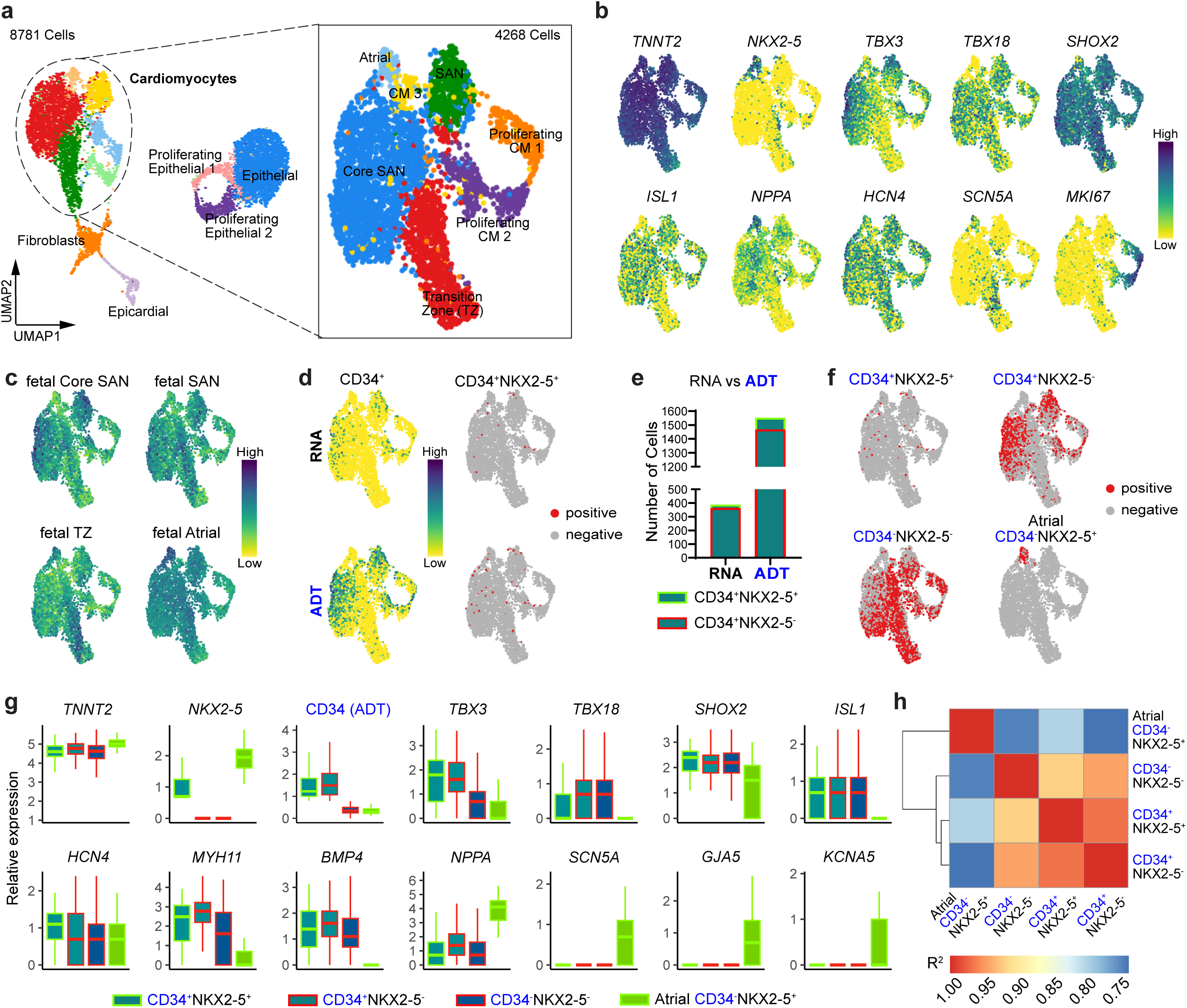
CITE-seq of hPSC-derived SANLPCs shows that CD34^+^NKX2-5^+^ and CD34^-^NKX2-5^-^ cardiomyocytes are transcriptionally similar to SAN pacemaker cells. **a** UMAP of the day 25 HES2-derived SANLPC CITE-seq dataset showing 11 cell clusters (left). Subclustering of the *TNNT2*^+^ cardiomyocytes showing 7 sub-clusters (right). **b** UMAPs of the subclustered cardiomyocytes showing the expression of the indicated genes. **c** UMAPs showing signature score distribution for the DEGs of the indicated fetal SAN cell types. **d** UMAPs showing the expression of CD34 on the transcript (RNA) or protein level (antibody-derived tag (ADT)) (left) and co-expression with NKX2-5 transcript (right). **e** Stacked bar graph quantifying the number of cells expressing CD34 by transcript (RNA) or protein (ADT) grouped based on NKX2-5 expression by transcript. **f** UMAPs showing the distribution of the selected cell populations indicated above. **g** Box plots showing the expression of the indicated genes on the transcript level and CD34 on the protein level (ADT) in the selected cell populations indicated below. **h** Spearman correlation between the selected cell populations indicated on the axes. p < 0.05 for all correlations. Blue labels indicate the data that is showing protein level (ADT) based CD34 expression.

We next analyzed the expression of CD34 at the transcript (RNA) and protein (antibody-derived tag, ADT) levels. A larger number of CD34^+^ cells was detected by ADT counts compared to transcript counts (1552 vs. 387) (Fig. 6d, e). Accordingly, we also identified a larger number of CD34^+^NKX2-5^+^ cells (85 vs. 24) when using the antibody-based detection of CD34^+^ cells. To further assess the identity of the different cell populations, we isolated CD34^+^NKX2-5^+^, CD34^-^NKX2-5^-^, and CD34^+^NKX2-5^-^ cells based on their expression of CD34 (ADT) and NKX2-5 (RNA) (Fig. 6f, Supplementary Fig. 9c). The distribution of NKX2-5^+^CD34^+^ and CD34^-^NKX2-5^-^ cells was scattered and did not line up with any specific cell cluster. In contrast, CD34^+^NKX2-5^-^ cells clearly segregate into the Core SAN and SAN cell clusters.

To characterize the CD34^+^NKX2-5^+^ and CD34^-^NKX2-5^-^ cells we compared the expression of typical pacemaker and working myocyte markers in these cells with the CD34^+^NKX2-5^-^ SAN pacemaker cells. We included CD34^-^NKX2-5^+^ myocytes from the Atrial cluster as an additional expression reference in this analysis. Both the CD34^+^NKX2-5^+^ and CD34^-^NKX2-5^-^ expressed comparable levels of the SAN pacemaker genes *TBX18, SHOX2, ISL1, HCN4, MYH11* and *BMP4* like the CD34^+^NKX2-5^-^ cells (Fig. 6g). Lower expression of these SAN genes was detected in the Atrial myocytes. Interestingly, expression of the pan-pacemaker marker *TBX3* was high in the CD34^+^NKX2-5^+^ and CD34^+^NKX2-5^-^cells, but lower in the CD34^-^NKX2-5^-^ cells. Expression of the atrial genes *NPPA*, *SCN5A*, *GJA5* and *KCNA5* was comparable between the CD34^+^NKX2-5^+^, CD34^-^ NKX2-5^-^, and CD34^+^NKX2-5^-^ cells and lower than in the Atrial myocytes. In addition, we performed Spearman correlation analysis using the DEGs of each of these cell populations, which showed that the CD34^+^NKX2-5^+^ and CD34^-^NKX2-5^-^ cells most closely correlate with the CD34^+^NKX2-5^-^ SAN pacemaker cells but not with the CD34^-^NKX2-5^+^ Atrial myocytes (Fig. 6h).

Collectively, these data indicate that the CD34^+^NKX2-5^+^ and CD34^-^NKX2-5^-^ cells are transcriptionally comparable to the CD34^+^NKX2-5^-^ SAN pacemaker cells and have a SAN pacemaker phenotype. The lower expression of *TBX3* in the CD34^-^NKX2-5^-^ cells suggests that they might be a SAN pacemaker progenitor population. This is consistent with the observation that the majority of the CD34^-^ sorted NKX2-5^-^ cardiomyocytes upregulate CD34 expression 25 days post-sort (Supplementary Fig. 7 f, g).

### CD34^+^ cardiomyocytes have a functional pacemaker phenotype

To assess if the hPSC-derived CD34^+^ cardiomyocytes also show the electrophysiological characteristics of pacemaker cells we recorded optical action potentials (Fig. 7a). SIRPA^+^CD90^-^NKX2-5^-^ FACS sorted SANLPCs, which we have previously shown to functionally represent pacemaker cells, were included as positive control^14^. In addition, ALCMs and VLCMs were included as reference cell types. We recorded fast spontaneous beating rates in MACS sorted CD34^+^ cells and SANLPC controls that were significantly faster than beating rates of VLCMs^14, 17^ (Fig. 7b). As expected for pacemaker cells, CD34^+^ cells and SANLPC controls displayed significantly slower maximum action potential upstroke velocities than ALCMs and VLCMs^14, 33^. Additional parameters that distinguish pacemaker cells from working cardiomyocytes are their action potential durations^14, 26, 33^. Both CD34^+^ cells and SANLPC controls showed significantly shorter action potential durations at 30% (APD30) and 90% (APD90) repolarization compared to VLCMs. The APD30/90 ratios for CD34^+^ cells and SANLPC controls were significantly larger compared to ALCMs which clearly distinguishes them from ALCMs. Finally, we analyzed the diastolic depolarization during phase 4 of the action potential, which is driven by funny currents specifically present in pacemaker cells^33,34^. Accordingly, we observed pronounced diastolic depolarization in CD34^+^ cells and SANLPC controls that was significantly faster than in ALCMs and VLCMs.

**Figure 7:**
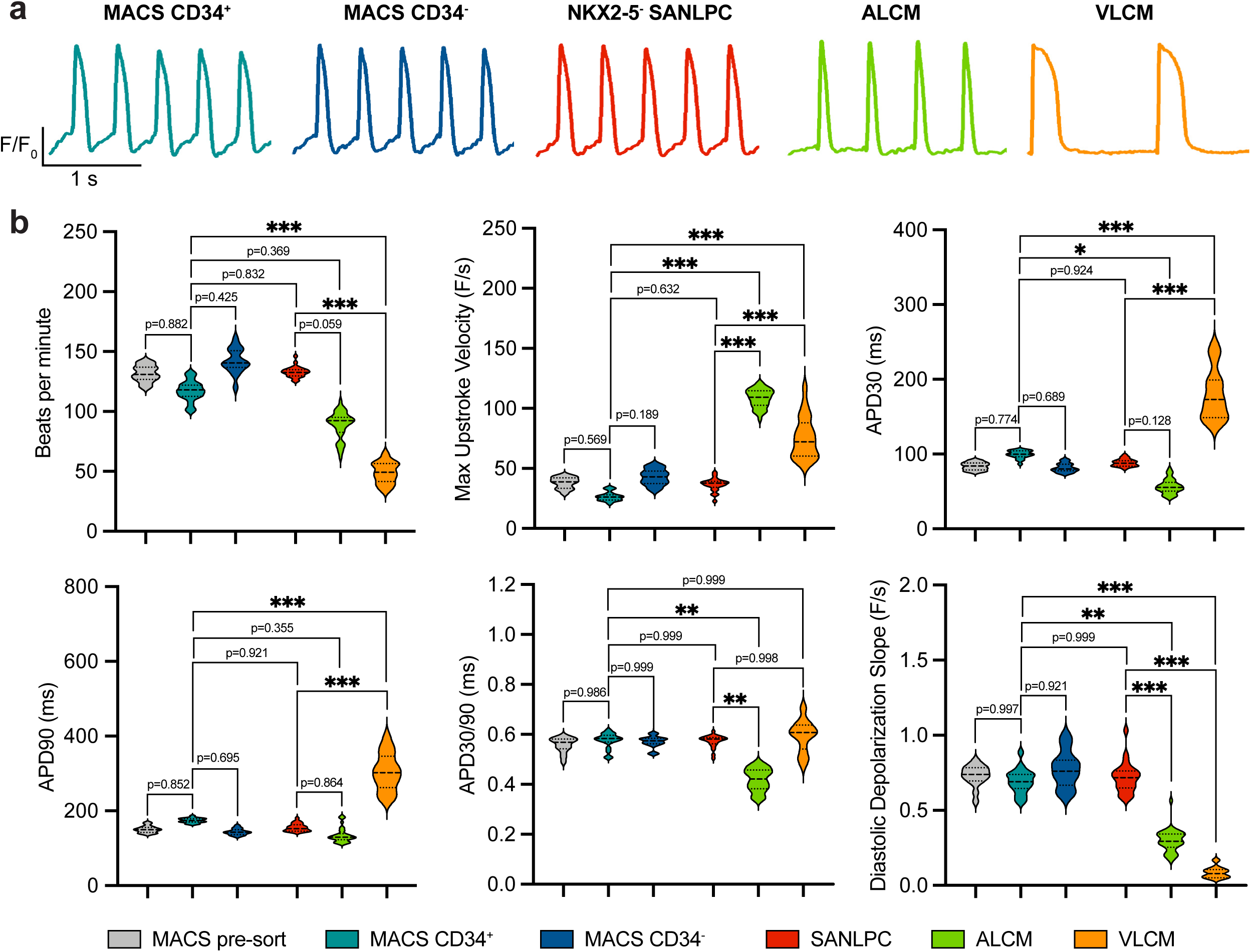
Optical action potential recordings confirm functional pacemaker phenotype of CD34^+^ sorted cells. **a** Representative optical action potential traces of day 25 MACS sorted CD34^+^ and CD34^-^ cells, FACS sorted SIRPA^+^CD90^-^NKX2-5^-^ SANLPCs, as well as ALCMs and VLCMs generated from the HES3 *NKX2-5^eGFP/w^* hPSC line. **b** Violin plots showing the optical action potential parameters analyzed in all five cell types (n ≥ 40 recordings from a total of three biological replicates (independent differentiations) for each cell type). Statistical analysis was performed using nested one-way ANOVA followed by Tukey’s post hoc test: *p < 0.05, **p < 0.01, ***p < 0.001 vs. indicated sample. APD30/90, Action potential duration at 30%/90% repolarization.

Interestingly, we did not observe significant functional differences between the CD34^+^ and CD34^-^ MACS sorted cells. This could be due to the fact that the CD34^-^ sorted cells still contained a sizable fraction of NKX2-5^-^ pacemaker cells (41 ± 3%, Fig. 5h, i), in addition to the NKX2-5^+^ Atrial and Transition Zone cells. In summary, our analysis of multiple action potential parameters clearly distinguished CD34^+^ cells from ALCMs and VLCMs and clearly showed that they have a functional pacemaker phenotype comparable to NKX2-5^-^ SANLPCs. These data therefore further confirm that CD34 is a reliable surface marker to isolate SAN pacemaker cells from hPSC-differentiation cultures.

## Discussion

The SAN is the primary pacemaker of the heart that initiates every single heartbeat. SAN failure is treated with the implantation of an electronic pacemaker device, which is associated with multiple downsides. Despite the important function of the SAN, its cellular and molecular composition is not completely resolved. This limits our ability to develop novel treatment options for SAN diseases. To date, no surface markers for SAN pacemaker cells have been identified, which represents a major bottleneck for disease modelling and the advancement of biological pacemaker cell therapy approaches. To address this and expand our knowledge especially on the cellular and molecular make-up of the human SAN, we performed snRNA-seq of human fetal SAN tissue and scRNA-seq of hPSC-derived SANLPCs. Our findings establish that the fetal SAN is composed of three distinct subtypes of pacemaker cells, including Core SAN, SAN, and Transitional Cells. Notably, the hPSC-derived cells closely resemble these fetal SAN pacemaker cells. Combining the fetal and hPSC-derived datasets allowed us to establish a shared list of Core SAN-specific genes. This list contains a host of novel markers, most importantly, the cell surface antigen CD34. We provide evidence that CD34 is specifically expressed by SAN pacemaker cells and that sorting for CD34^+^ cells from hPSC differentiations enriches for SANLPCs with a functional pacemaker phenotype.

Previous studies of the human SAN describe a core pacemaker structure that expresses *TBX3, TBX18, ISL1, HCN4,* and lacks the expression of *NKX2-5*^3–6, 12^. In addition, a transitional structure, also known as the paranodal area, located at the border zone between the SAN and atrial tissues, has been described. This transitional structure expresses a mix of pacemaker and atrial myocyte genes. Consistent with this, we identified Core SAN pacemaker cells and Transition Zone cells in our fetal snRNA-seq dataset. We also detected another pacemaker myocyte cell cluster that expressed lower levels of *TBX3* compared to the Core SAN cells that we annotated as SAN cells. Based on their expression pattern these cells most likely resemble the TBX18^+^, Shox2^+^, ISL1^+^ TBX3^low^ and NKX2-5^-^ sinus venosus myocardium of the developing heart^2, 21, 35, 36^. Analysis of hPSC-derived SAN differentiation cultures also identified Core SAN, SAN and Transition Zone cells. Comparison of the hPSC-derived cells to the fetal counterparts showed a striking molecular overlap. This finding strongly supports the concept that the *in vitro* generated cells are indeed SAN pacemaker cardiomyocytes.

Studies of the SAN in the mouse heart identified a NKX2-5^-^TBX3^+^TBX18^+^ SAN head, located at the boundary between the superior vena cava and the right atria, and a NKX2- 5^+^TBX3^+^TBX18^-^ SAN tail that extends along the terminal crest towards the inferior vena cava^7, 9, 37–40^. In the human heart, a SAN head and tail can be anatomically defined^6, 41^. However, a clear molecular distinction between a SAN head and SAN tail, like in the mouse, has not been shown. In agreement with that, we did not identify a SAN tail cell population in our fetal nor hPSC-derived datasets. Together our molecular analysis provides detailed insights into the composition of the developing human SAN. In addition, our study corroborates that hPSC differentiations can replicate SAN development including the development of the different subtypes of pacemaker cells.

To identify markers that can be used *in vivo* and *in vitro* to detect, target, and isolate the rare pacemaker cells of the Core SAN, we combined the fetal snRNA-seq and hPSC-derived scRNA-seq datasets. This analysis resulted in a list of 36 genes that contain an abundance of new markers. Amongst the most highly ranked markers was *MYH11,* the gene encoding for the smooth muscle-specific myosin heavy chain 11. *MYH11* is expressed in smooth muscle cells of the digestive system, bladder, aorta, and coronary arteries of the heart. Previous transcriptomic studies have detected *MYH11* in SAN myocytes^4, 8, 10, 18, 21^. Our study extended these findings to show that MYH11 protein is specifically expressed by hPSC-derived SANLPCs and fetal SAN pacemaker cells but not AVN, atrial, nor ventricular cardiomyocytes. We and others also found that additional smooth muscle genes (*ACTA2, MYLK*) are specifically expressed in SAN myocytes^4, 8, 10, 18^. It remains to be explored whether these genes play an important role in SAN physiology. As a second gene from our Core SAN marker list, we validated the expression of the TGF-β signaling ligand BMP4 specifically in SAN pacemaker cardiomyocytes. BMP signaling has previously been shown to play an important role during SAN development in both animal models as well as hPSC- differentiations, where BMP4 is used to induce the specification of SAN pacemaker cells from cardiogenic mesoderm^4, 8, 14, 21, 42^. The marker list further contained several genes involved in calcium binding and calcium channel regulation (*ANXA2, NECAB1, SPOCK1, RRAD, PRKG1*). This is consistent with the well-established role of calcium ions in the regulation of the spontaneous action potential firing of the SAN (Ca^2+^-clock)^43^. In addition, we identified several genes commonly expressed in neurons as Core SAN markers (*BASP1, PPFIA2, EFR3B, SLIT2, LYPD6*). These genes play a role in axon growth and guidance which is in keeping with the finding that Neuron Projection Guidance/Development were identified as top GO terms for the fetal/hPSC-derived SAN cells. The SAN tissue is highly innervated by the autonomous nervous system which led to the detection of neuronal genes in past bulk RNA-seq experiments^4, 8^. Here we analyzed gene expression on the single-cell level within *TNNT2*^+^ cardiomyocytes making it highly unlikely that gene expression from neuronal cells was detected. Similarly, multiple neuronal genes were detected in SAN pacemaker cells in recent sc/snRNA-Seq studies of the mouse and human SAN^9, 10, 12, 44^. Collectively, these findings suggest that SAN pacemaker cells express a subset of neuronal genes, potentially utilize comparable transcriptional and signaling networks as neurons, and actively participate in guiding the axons of the innervating autonomic nervous system. These are interesting concepts that need to be further explored in the future.

In addition to the above set of genes we also identified a number of cell surface markers that were specifically expressed in Core SAN pacemaker cells, including *CD34*, *CADM1*, *EFNB2*, *NTRK2*, and *ELAPOR2*. We focused our analysis on the single pass transmembrane glycoprotein CD34, because of its routine use as surface marker for cell isolations. Classically, CD34 is known as a selection marker for hematopoietic stem and progenitor cells, as well as for its expression in endothelial cells^45–47^. In recent years, CD34 has also been described to mark tissue resident progenitors such as muscle satellite cells and mesenchymal stem cells^48, 49^. Additionally, there have been reports of CD34 expression in telocytes, an interstitial cell type that can be found throughout the body, including the heart^28, 50^. Here we extend these findings and show expression of CD34 in SAN pacemaker myocytes of the human heart and in hPSC-derived SAN pacemaker cells. Although CD34 is a well-established marker, especially in the hematopoietic system, its functional role is not well established. One of CD34’s potential functions relates to cell migration and adhesion via binding of selectins by its extracellular mucin domain^51^. Further studies are needed to address whether CD34 expression on developing SAN cells impacts their migratory behavior and overall SAN tissue organization. Interestingly, multiple genome-wide association studies have implicated genetic variants in the CD34 gene in heart rate changes, indicating that CD34 might have a physiological role beyond cell migration in SAN cells^52–54^.

We used FACS and MACS sorting for CD34^+^ cells to demonstrate that this approach enables transgene independent and straightforward isolation of SAN-like pacemaker cells from multiple hPSC lines. Our analysis showed that these CD34^+^ sorted cells display molecular and functional properties of SAN pacemaker cells. This novel SAN cell surface marker will be highly valuable for future SAN specific disease modelling and drug screening projects using patient-derived iPSCs, to gain a better understanding of the mechanisms of sinus node diseases and to develop novel therapies. It will further enable the enrichment of SAN pacemaker cells for toxicology and safety pharmacology screens. From a cell therapy perspective, MACS will allow to efficiently isolate the large numbers of CD34^+^ SAN pacemaker cells from hPSC-differentiation cultures that will be required for future biological pacemaker applications^14, 55^. Conversely, MACS-based depletion of CD34^+^ cells can be used to remove unwanted pacemaker cells from populations of hPSC-derived ventricular cardiomyocytes. This could reduce the risk of unwanted arrhythmias in cell replacement therapies for patients who suffered a myocardial infarction^56, 57^. Beyond that, CD34 specific antibodies conjugated to a drug or other therapeutic cargo could be used in the future to specifically deliver therapeutics to the SAN in patients with too slow or too fast sinus rhythm^25, 58^. To avoid cross reaction to other CD34^+^ cells in the body, bispecific antibodies that target both CD34 and a myocyte specific epitope such as SIRPA or HCN4 could be employed^59^.

When we compared our human Core SAN marker list to previously published mouse scRNA-seq data^9, 25^, we found that only half of the markers were conserved. Specifically, CD34 expression was not detected in mouse SAN pacemaker cells. Likewise, the majority of the previously reported mouse SAN marker genes were not specifically expressed by human SAN pacemaker cells. These species differences are important when choosing markers for the characterization and isolation of hPSC-derived SAN-like pacemaker cells as well as for the clinical translation of SAN cell surface marker-based therapeutics. Hence, the insights that our study of the human SAN provides will be a valuable resource for the scientific community.

In summary, our study provides the first transcriptomic profile of the developing human SAN at single-cell resolution. Using this data, we showed that hPSC-derived SANLPCs closely resemble the SAN pacemaker cells in the fetal heart and identified a shared set of 36 markers for the identification of SAN pacemaker cells *in vitro* and *in vivo*. This list contains a host of new markers most notably CD34, a novel SAN cell surface marker. We show that CD34 is specifically expressed by human fetal SAN pacemaker myocytes and that it can be used to isolate SANLPCs from hPSC differentiation cultures. Our findings have important implications for future iPSC-based SAN disease modelling studies as well as cell therapy approaches using hPSC-derived biological pacemakers. The markers identified in our study will also advance our ability to specifically deliver therapeutics to SAN cells *in vivo*, using antibody-based approaches. Taken together, our findings will contribute to a better understanding of SAN diseases and to improved future treatment options for patients.

## Methods

### Mice

Wild-type, P0-3 CD1 mice were obtained from the Animal Resource Centre at UHN. All animal work was performed in accordance with the Animal Use and Care Committee at UHN. Both female and male mice were used.

### Human fetal heart samples

Human fetal heart samples gestation week 17-20 were obtained from the Mount Sinai Hospital Research Centre for Women’s and Infants Health BioBank, Toronto. Work with the human fetal tissue was approved by the Research Ethics Boards of the Mount Sinai Hospital, Toronto and the University Health Network (UHN), Toronto. Fetuses with chromosomal abnormalities, cardiovascular defects, congenital heart defects, and arrhythmias were excluded from this study. The fetal sex was not specified for this study and not included in the health information shared with the study team.

### hPSC lines

Human pluripotent stem cell lines HES3-Nkx2-5^egfp/w^ ^30^, and HES2^60^ were used in this study. Work with hPSCs was approved by the Stem Cell Oversight Committee of the Canadian Institutes of Health Research.

### Maintenance and directed differentiation of hPSCs

The human pluripotent stem cell lines were maintained on irradiated mouse embryonic fibroblasts in DMEMF/12 (ThermoFisher, MT-10-092- CV) supplemented with 20% KnockOut serum replacement (ThermoFisher, 10828028), 2mM L-glutamine (ThermoFisher, 25030081), 1x non-essential amino acids (ThermoFisher, 11140050), 55μM β-Mercaptoethanol (ThermoFisher, 21985023) and 1% penicillin/streptomycin (ThermoFisher, 15070-063), and 20ng/ml recombinant human (rh)-bFGF (Biotechne, 233-FB/CF) as described previously^61^. All cell lines used in this study had a normal karyotype and tested negative for mycoplasma contamination.

For directed differentiations into the different cardiomyocyte subtypes we used our previously reported embryoid body (EB)-based protocols with the following modifications^14, 26^. StemPro-34 Media (ThermoFisher, 10639011) supplemented with 1% penicillin/streptomycin, 2mM L-glutamine, 50μg/mL ascorbic acid (MilliporeSigma, A4544), 75μg/mL transferrin (MilliporeSigma, 10652202001) and 50μg/mL monothioglycerol (MilliporeSigma, M6145) was used as a base media for all differentiation steps. hPSCs grown to ∼80% confluence were dissociated into single cells using TrypLE (ThermoFisher, 12605010). Cells were suspended in base media containing 20% StemPro-34 diluted with IMDM (ThermoFisher, 12440061) supplemented with 0.01x ITSX (ThermoFisher, 51500056), 1ng/mL rhBMP4 (Biotechne, 314-BP/CF), 10μM ROCK inhibitor Y27632 (Selleckchem, S1049), and 250U/ml DNase I (MilliporeSigma, 260913) at a concentration of 1×10^6^ cells/mL and cultured for 18h on an orbital shaker at 80rpm in low attachment 6-cm dishes for the formation of EBs. EBs were cultured in a low oxygen environment (5% CO_2_, 5% O_2_, 90% N_2_) until day 11 and then in a 5% CO_2_ ambient air environment for the remainder of the culture period. Tissue culture plastic was pre-coated with 5% (w/v) poly-HEMA (MilliporeSigma, P3932) to generate low adherence plates for the maintenance of EBs in suspension culture.

#### Ventricular-like cardiomyocyte (VLCM) differentiation

At day 1 of differentiation the EBs were moved into ventricular mesoderm induction media consisting of base media containing 100% StempPro-34 supplemented with 1μM CHIR99021 (Biotechne, 4423), 2.5ng/ml rhbFGF, 10ng/ml rhBMP4, and 8ng/ml rhActivinA (Biotechne, 338-AC/CF). At day 3 the EBs were washed once with IMDM and transferred to cardiac induction media that consisted of base media containing 100% StemPro-34 supplemented with 1uM IWP2 (Biotechne, 3533) and 10ng/ml rhVEGF (Biotechne, 293-VE/CF). At day 5 media was changed to cardiac maintenance media that consisted of base media containing 20% StemPro-34 diluted with IMDM supplemented with 5ng/ml rhVEGF. Media was changed every 3-4 days and starting at day 11 EBs were cultured in maintenance media without rhVEGF.

#### Atrial-like cardiomyocyte (ALCM) differentiation

At day 1 of differentiation the EBs were moved into atrial mesoderm induction media consisting of base media containing 100% StempPro-34 supplemented with 1μM CHIR99021 (Biotechne, 4423), 2.5ng/ml rhbFGF, 4ng/ml rhBMP4, and 3ng/ml rhActivinA. At day 3 the EBs were washed once with IMDM and transferred to cardiac induction media as above (VLCM differentiation) that was additionally supplemented with 0.5uM retinoic acid (MilliporeSigma, R2625). From day 5 onwards the differentiation was continued as described for VLCMs.

#### Sinoatrial Node-like pacemaker cell (SANLPC) differentiation

At day 1 of differentiation the EBs were moved into pacemaker mesoderm induction media consisting of base media containing 100% StempPro-34 supplemented with 1μM CHIR99021 (Biotechne, 4423), 2.5ng/ml rhbFGF, 4ng/ml rhBMP4, and 3ng/ml rhActivinA. At day 3 the EBs were washed once with IMDM and transferred to pacemaker induction media that consisted of base media containing 100% StemPro-34 supplemented with 0.5uM IWP2 (Biotechne, 3533), 10ng/ml rhVEGF, 2.5ng/ml rhBMP4, 1.5uM SB-431542 (MilliporeSigma, S4317), and 0.125uM retinoic acid. 500-1000nM PD173074 (Biotechne, 3044) were added to the media on either day 3 (HES3-Nkx2-5^egfp/w^ line) or day 4 (HES2 cell line). At day 6 media was changed to base media containing 100% StemPro-34 supplemented with 5ng/ml rhVEGF. At day 8 media was changed to cardiac maintenance media and differentiations were continued as described above for VLCMs.

### Flow cytometry and cell sorting

Day 6-8 EBs were dissociated using TrypLE for 3-6 minutes at 37°C. Day 10-50 EBs were dissociated using Collagenase type 2 (1 mgl/ml, Worthington, 4176) in HANKs buffer for 16 hours at room temperature, followed by treatment with TrypLE as above if further dissociation was required. The cells were stained using the following antibodies: anti- CD172a/b-PE-Cy7 (SIRPa) (Biolegend, 323808, 1:1000), anti-CD90-BV421 (BD, 562556, 1:300), anti-CD31-PE (BD, 555446, 1:50), anti-CD34-APC (BD, 340441, 1:300), anti-CD34-PE-Cy7 (ThermoFisher, 25-0349-42, 1:100), anti-Cardiac Troponin T (cTNT) (ThermoFisher, MA5-12960, 1:2000), anti-NKX2-5 (Cell Signaling, 8792S, 1:1000). To detect unconjugated primary antibodies the following secondary antibodies were used: goat anti-mouse IgG-PE (Jackson ImmunoResearch, 115-115-164; 1:500), donkey anti-rabbit IgG-A647 (ThermoFisher, A31573; 1:500)

For cell-surface markers, cells were stained PBS containing 5% FCS and 0.02% sodium azide for 30 minutes at 4°C. For intracellular staining (cTNT, NKX2-5), cells were fixed using 4% PFA for 10 minutes at 4°C and stained in PBS containing 0.3% BSA, 0.3% TritonX over night at 4°C. Stained cells were analyzed using the LSRFortessa flow cytometer (BD) or Cytoflex flow cytometer (Beckman Coulter). For FACS, the cells were kept in IMDM containing 0.5% FCS and sorted at a concentration of 10 million cells/ml using a FACSAria (BD) sorter at the SickKids-UHN Flow Cytometer Facility. For MACS, the PSC-Derived Cardiomyocyte Isolation Kit (Miltenyi, 130-110- 188) and the CD34 MicroBead Kit (Miltenyi, 130-046-702) were used according to the manufacturer’s instructions. All data was analyzed using FlowJo software (BD). The gating strategies used for data analysis are shown in Supplementary Fig. 10.

### Immunohistochemistry

For cultured cells, day25 EBs were dissociated, and MACS sorted using the PSC-Derived Cardiomyocyte Isolation Kit to enrich for cardiomyocytes as described above. Cells were plated on 12mm cover glasses (FisherScientific, 22293232P) precoated with Matrigel (25%, FisherScientific, CB40230) and cultured for 3-5 days until confluent monolayers were obtained. Cells were fixed with 4% PFA in PBS for 10 minutes at 4°C and permeabilized using PBS containing 0.3% TritonX, 200 mM glycerin (MilliporeSigma, G2289) for 20 min at room temperature (RT). Samples were blocked with PBS containing 10% donkey serum, 2% BSA, and 0.1% TritonX (blocking buffer) for 30 minutes at RT before incubating with primary antibodies in blocking buffer overnight at 4°C. The samples were washed three times with PBS containing 0.1% BSA, 0.1% TritonX (wash buffer), before applying secondary antibodies together with 10ug/ml Hoechst (FisherScientific, H3570) in blocking buffer for 1 hour at RT. The samples were washed three times with wash buffer before mounting with ProLong Diamond Antifade Mounting Media (FisherScientific, P36965).

To prepare tissue sections of human gestation week 17-20 SAN and AVN we followed a previously described dissection protocol (Supplementary Fig. 5a) ^27^. Briefly, the hearts were placed with the anterior side facing up and the ventricular chambers were dissected away leaving a small rim of ventricular tissue apical to the mitral and tricuspid valves. Next, the right atria was opened by cutting through the tricuspid valve towards the superior vena cava on the anterior side of the heart. Next the atrial free wall was cut open, flattened, and pinned down revealing the view of the crista terminalis, the SAN and the AVN. Sections were prepared from the posterior (epicedial) side of the dissected tissue. To obtain tissue sections of P0-3 mouse SAN whole hearts were isolated, and cross sections were prepared from the base of the hearts.

All tissue preparations were fixed overnight in 4% formalin, transferred to 70% ethanol, and sent for paraffin embedding and preparation of 5um thick sections at the Pathology Research Program at UHN. Sections were re-hydrated using decreasing xylene/ethanol solutions before heat-induced antigen retrieval in citrate buffer (pH 6). Sections were blocked in PBS containing 0.5% donkey serum and incubated with primary antibodies overnight at 4°C. The samples were washed three times with PBS before applying secondary antibodies together with 10ug/ml DAPI (FisherScientific, D1306) in blocking buffer for 1 hour at RT. The samples were washed three times with PBS and once with distilled water before mounting with ProLong Diamond Antifade Mounting Media.

The following primary antibodies were used: anti-BMP4 (Abcam, ab124715, 1:50), anti-Cardiac Troponin T (cTNT)-FITC (Miltenyi, 130-119-575, 1:100), anti-Cardiac Troponin T (cTNT) (ThermoFisher, MA5-12960, 1:200), anti-CD31 (DAKO, M0823, 1:50), anti-CD34 (Abcam, ab81289, 1:300), anti-CD34 (BD, 340441 1:100), anti-GFP (Abcam, ab13970, 1:500), anti-HCN4 (Antibodies Inc, 75-150, 1:100), anti-MLC2V (Abcam, ab79935, 1:100), anti-MSX2 (Abcam, ab223692, 1:300), anti-MYH11 (Abcam, ab224804, 1:100), anti-NKX2-5 (Cell Signaling, 8792S, 1:1000), anti-NPPA (Abcam, ab209232, 1:300), anti-SHOX2 (Abcam, ab55740, 1:300), and anti- TBX3 (ThermoFisher, 424800, 1:100). The following secondary antibodies were used: Donkey anti-rabbit IgG-A647 (ThermoFisher, A31573, 1:500), Donkey anti-rabbit IgG-A555 (ThermoFisher, A31572, 1:500), Donkey anti-mouse IgG-A647 (ThermoFisher, A31571, 1:500), Donkey anti-mouse IgG-A555 (ThermoFisher, A31570, 1:500), and Donkey anti-chicken IgY-A488 (ThermoFisher, A78948, 1:500). Masson’s Trichrome staining was performed by the UHN Pathology Research Program.

All samples were imaged at UHN’s Advanced Optical Microscopy Facility using the Nikon A1R confocal microscope and associated NIS-Elements software (Nikon). Whole-slide images were scanned using the 20X objective lens on the Scan-scope AT2 brightfield scanner (Aperio). Images were processed and prepared using ImageJ (FIJI)^62^.

### Optical action potential recordings

All cell types analyzed by optical action potential recording were generated from the HES3 *NKX2-5^eGFP/w^* hPSC line. Day 25 EBs were dissociated into single cells as described above and re-aggregated into smaller aggregates (∼50-100μm diameter) by plating into 96 well ultra-low attachment plates (FisherScientific, 07-200-603) at 200,000 cells/well. To obtain enriched populations of SANLPCs SIRPA^+^CD90^-^NKX2-5^-^ cells were isolated by FACS. CD34^+^ and CD34^-^ cells were isolated from SANLPC differentiation cultures by MACS. Cells were allowed to recover for 3-5 days and stained at 37°C, 5% CO_2_ in salt-adjusted IMDM (final concentrations in mmol/l: 137 NaCl, 4.4 KCl, 1.5 CaCl_2_, 25 HEPES, and 25 D-glucose) (ThermoFisher, 12440061) using the FluoVolt Membrane Potential Kit (ThermoFisher, F10488) according to the manufacturer’s instructions. Imaging of optical action potentials was performed in salt-adjusted IMDM at 37°C using a pco.edge 4.2Q sCMOS camera mounted on a Nikon Ti2-E inverted microscope with a CFI60 10X objective. Recordings of at least 10 seconds were acquired at 400 frames/second using NIS-Elements software (Nikon). Optical action potential characteristics (frequency, maximal upstroke velocity, and action potential duration) were analyzed using custom Python scripts and standard techniques previously desecribed^63–65^. Briefly, signals were pre-processed to correct for staining related artifacts, and filtered to remove high-frequency noise, using a low-pass Butterworth filter with a cut-off frequency of 100Hz. Recordings were segmented into individual action potentials that were then normalized in intensity and aligned for ensemble averaging (n ≥ 5 action potentials were averaged per recording). To determine the diastolic depolarization, the maxima of the second derivative of an over-filtered representation of the original signal was used, to detect the point of transition from the action potential phase 4 depolarization to action potential phase 0 depolarization. The phase 4 portion of the action potential was extracted, and linear regression analysis was used to determine the slope of depolarization.

### Quantitative reverse transcription PCR

Total RNA from hPSC-derived cardiomyocytes was isolated using the RNAqueous-Micro Total RNA isolation kit including RNase-free DNase treatment (ThermoFisher, AM1931). cDNA was prepared by reverse transcribing the isolated RNA using oligo(dT) primers and random hexamers with Superscript III Reverse Transcriptase (ThermoFisher, 18080044). RT-qPCRs were performed using the QuantiNova SYBR Green PCR kit (Qiagen, 208057) according to the manufacturer’s instructions on the QuantStudio5 RT-qPCR machine (ThermoFisher). Each experiment included a tenfold dilution series ranging from 25ng/mL to 2.5pg/mL of human genomic DNA standards to evaluate PCR efficiency and to calculate the copy number of each gene relative to the house keeping gene TBP as described previously^31^. Primer sequences are listed in Supplementary Table 5.

### Sample processing for single-nucleus RNA sequencing

Single-nucleus RNA-seq was performed on a gestation week 19 fetal heart. To capture the tissue containing the SAN, we dissected the heart and isolated the tissue at the boundary of the right atria with the superior vena cava (Fig. 1a). The tissue was immediately frozen using liquid nitrogen and stored at -80°C. The frozen tissue was retrieved and dissociated using lysis buffer containing: 0.32M sucrose (Sigma, S0389), 5mM CaCl_2_ (Fluka, 21114), 3mM Mg(AC)_2_ (Sigma, 228648), 20mM Tris-HCl (ThermoFisher, 15567-027), 0.1%Triton X-100 (MilliporeSigma, X100), 0.1mM EDTA (ThermoFisher, AM9260G) in DNase/RNase free water (Qiagen, 129115). A douncer was used to mechanically homogenize the tissue until single nuclei were obtained. Homogenate was washed twice with PBS containing 1% BSA and 0.2U/ul RNAseOUT (ThermoFisher, 10777019) and filtered through a 40um cell strainer. Nuclei were stained with DAPI (10μg/ml) and sorted for DAPI^+^ single-nuclei on a FACSAria (BD) sorter at the SickKids-UHN Flow Cytometer Facility.

### Sample processing for single-cell RNA sequencing and CITE sequencing

Single-cell RNA-seq was performed on day 25 SANLPC differentiation cultures of the HES2 cell line (Fig. 2a). The EB’s were dissociated and stained with DAPI as described above before being sorted for DAPI^-^ single live cells using a FACSAria (BD) sorter at the SickKids-UHN Flow Cytometer Facility. For CITE-seq, day 25 SANLPC differentiation cultures of the HES2 cell line were processed as described above. Before sorting, samples were stained with DAPI and the hCD34 TotalSeq^TM^-B0054 antibody (Biolegend, 343539) and DAPI^-^ single cells were collected as detailed above. The optimal concentration of the hCD34 TotalSeq^TM^ antibody was determined using flow cytometric analysis with an anti-mouse AF647 secondary antibody (ThermoFisher, A-31571). The hCD34 TotalSeq^TM^ antibody concentration that matched to the proportion of CD34^+^ seen when using the anti-CD34 APC antibody (BD, 340441 1:300) was chosen for the CITE-Seq experiment (Supplementary Fig. 9a).

### Single-cell/nuclei RNA sequencing, raw data processing, quality control, clustering, and differential gene expression analysis

Single-cell and single-nuclei suspensions were processed for sequencing on the 10x Genomics platform using the Chromium Single Cell 3’ v3 reagent kit. Single-cell/nuclei libraries were sequenced on the Illumina NovaSeq 6000 with a sequencing depth of > 45,000 reads per cell. Raw data processing was performed with the 10x Genomics CellRanger pipeline by UHN’s Princess Margaret Genomics Centre. Single-cell RNA-seq data and single-nuclei RNA-seq data were mapped to the human reference Genome GRCh38 and GRCh38- premrna respectively. Downstream data analysis was performed in R using the Seurat toolkit (version 4.0.2, https://satijalab.org/seurat/)^66^. First, data was filtered to remove low quality cells and potential doublets by removing cells with low library size (<1500) and high library size (>50000). In addition, a cutoff for mitochondrial gene expression was applied to remove damaged cells (fetal snRNA seq: 1%, hPSC scRNA-seq: 31.512%, hPSC CITE-seq 21%). Data was normalized using ScTransform, principal components analysis was performed using RunPCA, and the top 25 principal components were used for clustering into distinct cell clusters (FindClusters function). The scClustViz tool^67^ was used to determine optimal cluster resolution by increasing resolution until the minimal number of differentially expressed genes between each cluster reached 0-10. The following final cluster resolutions were used in this study: fetal snRNA-seq: 0.8, hPSC scRNA-seq: 0.8, hPSC CITE-seq: 0.4. The FindAllMarkers function (only.pos = TRUE, min.pct = 0.1, logfc.threshold = 0.25) was used to identify differentially expressed genes (DEGs) within each cluster or between subsets of selected cells. The top DEGs were identified after sorting the gene lists based on average log-fold expression changes. The genes used to identify which cell type each cluster represents are listed in Supplementary Table 1. To generate datasets that only contain cardiomyocytes, cell clusters expressing *TNNT2* were selected for subclustering. The genes used to identify the cardiomyocyte subtype clusters are discussed in the main text. Uniform manifold approximation and projection (UMAP) was used for dimensionality reduction to visualize the data in two dimensions. To read out CD34 expression on the protein level in the CITE-seq dataset, raw counts from the antibody-derived tag (ADT) were normalized with NormalizeData function (method = CLR) and a cut off of <0.84 was set to match CD34 positive expression in flow cytometric data (Supplementary Fig. 9). For selection of cells by RNA expression, a cut off of >0 was used for both NKX2-5 and CD34.

### Gene signature score and gene ontology analysis

Gene signature scores were obtained using UCell^22^ for the top 200 positive differentially expressed genes from the comparison groups. Signature score distributions were visualized using UMAPs. Gene ontology analyses were performed using the Database for Annotation, Visualization, and Integrated Discovery (DAVID) Bioinformatics Resources (version Dec. 2021, https://david.ncifcrf.gov/summary.jsp). Analysis was focused on biological processes (GOTERM_BP_all).

### Harmony data integration of the fetal snRNA-seq and hPSC scRNA-seq datasets

The fetal and hPSCs datasets were normalized and clustered using Seurat. Each dataset was normalization using ScTransform. Next, both samples were scaled separately as it has been shown to facilitate integration of single-cell and single-nucleus RNA-seq datasets in prior studies^68^. The two datasets were then concatenated to generate a merged gene expression matrix. The merged dataset was scaled and principal component analysis was applied to the data for dimension reduction. We then applied Harmony (version 1.0) integration on the principal components to remove the technical batch variations. UMAP was next applied to Harmony-adjusted top components for visualization.

### Spearman correlation analysis

To compare the cell types in the fetal and hPSC datasets correlation analysis was performed. The list of DEGs for each fetal cluster was sorted based on fold change and the top 200 genes were selected as the feature set for the correlation analysis. We then restricted the comparison to cell clusters of interest from each dataset (fetal: Core SAN, SAN, Transition Zone, Atrial, Epithelial cells, Epicardial cells, Fibroblasts; hPSC: Core SAN, SAN, Transition Zone, Atrial, Epithelial cells, Epicardial cells, Fibroblasts). Duplicate features and the genes not detected in the hPSC sample were removed. The resulting genes were used to calculate Spearman correlation between the fetal and hPSC clusters (Fig. 2k). To compare the CD34^+^ NKX2-5^+^, CD34^+^NKX2-5^-^, CD34^-^NKX2-5^-^, and CD34^-^NKX2-5^+^ Atrial cells in the CITE-Seq dataset a similar correlation analysis was performed with the following modifications. The positive DEGs from each of the selected cell types were selected as the feature set for the correlation analysis. The resulting genes were used to calculate Spearman correlation between the selected cell types (Fig. 6h).

### Comparison of mouse and human datasets

The raw data deposited by Goodyer et al.^9^ for the single-cell RNA-seq dataset of the mouse SAN was downloaded and analyzed using Seurat with the same settings as described in the original publication. For the comparison of gene expression in mouse and human SAN cells the 36 Core SAN DEGs contained in both human fetal and hPSC datasets were selected. In addition, the top 25 SAN DEGs^9^ and the top 25 SAN surface marker DEGs^25^ from the mouse dataset were selected (sorted based on fold change). To visualize the gene expression in each dataset, bar graphs were generated in R. Genes from our human datasets were categorized as conserved if their expression in the mouse Core SAN cells was higher than in the Atrial cells. Genes from the mouse dataset were categorized as conserved if their expression in the human Core SAN cells was higher than in the Atrial cells in at least one of the human datasets, fetal or hPSC. If gene expression in the Core SAN was below 0.1 the gene was considered not expressed and listed as not conserved. These low expressing genes were also visually confirmed looking at the gene expression in UMAPs.

### Quantification and Statistical Analysis

All data are presented as mean ± standard error of the mean (SEM). Indicated sample sizes (n) represent biological replicates (cell culture replicates or primary tissue samples). Sample sizes from single-cell datasets represent the number of cells analyzed as indicated in the individual figures. Statistical significance was determined using GraphPad Prism 9 (version 9.5.1). For the comparison between two conditions Student’s t-test (unpaired, two-tailed) was applied. For the comparison between multiple conditions one-way ANOVA with Bonferroni post hoc test was applied. Results were considered significant at p < 0.05 (*), p < 0.01 (**), and p < 0.001 (***). All statistical parameters are detailed in the respective figures and figure legends. Due to the nature of the study no randomization or blinding was implemented and no statistical methods were used to pre-determine samples sizes.

### Figure preparation

Figures were prepared using Adobe Illustrator. Schematics in Fig. 1a and Fig. 2a were generated using Biorender: https://biorender.com.

### Data and Software availability

The data supporting the findings of this study are available within the article and the supplementary information files. Any additional information is available from the corresponding author upon request. Raw sc/snRNA-seq data generated in this study have been deposited at GEO under the following accession number: [will be added once data is deposited]. We did not generate any original code in this study and all applied software packages are detailed and referenced in the method section.

## Acknowledgements

We would like to thank the members of the Protze laboratory for experimental advice and critical comments on the manuscript. A. Elefanty and E. Stanley (Monash University, Victoria, AU) for providing the HES3 NKX2-5^egfp/w^ reporter line, the SickKids-UHN Flow Cytometry Facility for assistance with cell sorting, the Advanced Optical Microscopy Facility at UHN for assistance with confocal microscopy, the Princess Margaret Genomics Centre at UHN for assistance with sc/snRNA-seq and data processing, and the Mount Sinai Hospital Research Centre for Women’s and Infants Health BioBank for providing access to fetal heart samples. This work was supported by grants from the Canadian Stem Cell Network (IRP-ECI-19, Protze), the Canadian Institute of Health Research (CIHR PJT 169090), the Canadian Foundation for Innovation (John R. Evans Leaders Fund, 38651), and funding from BlueRock Therapeutics LP.

## Author contributions

A.A.L. designed the project, performed experiments, analyzed data, and wrote the manuscript. A.L., D.P., T.T., G.B., and S.P., generated and analyzed the sc/snRNA-seq data. M.A., K.H., and Z.L., designed, performed, and analyzed the optical action potential experiments. M.L., M.L.C., and B.M.M., performed and analyzed tissue culture, flow cytometry, and immunohistochemistry experiments and provided valuable input on the manuscript. A.M., assisted with the dissection of fetal heart tissues. S.P. designed the project and wrote the manuscript.

## Materials & Correspondence

Any request for material should be addressed to S.P.

---

**Supplementary Figure 1 (related to Figure 2):**
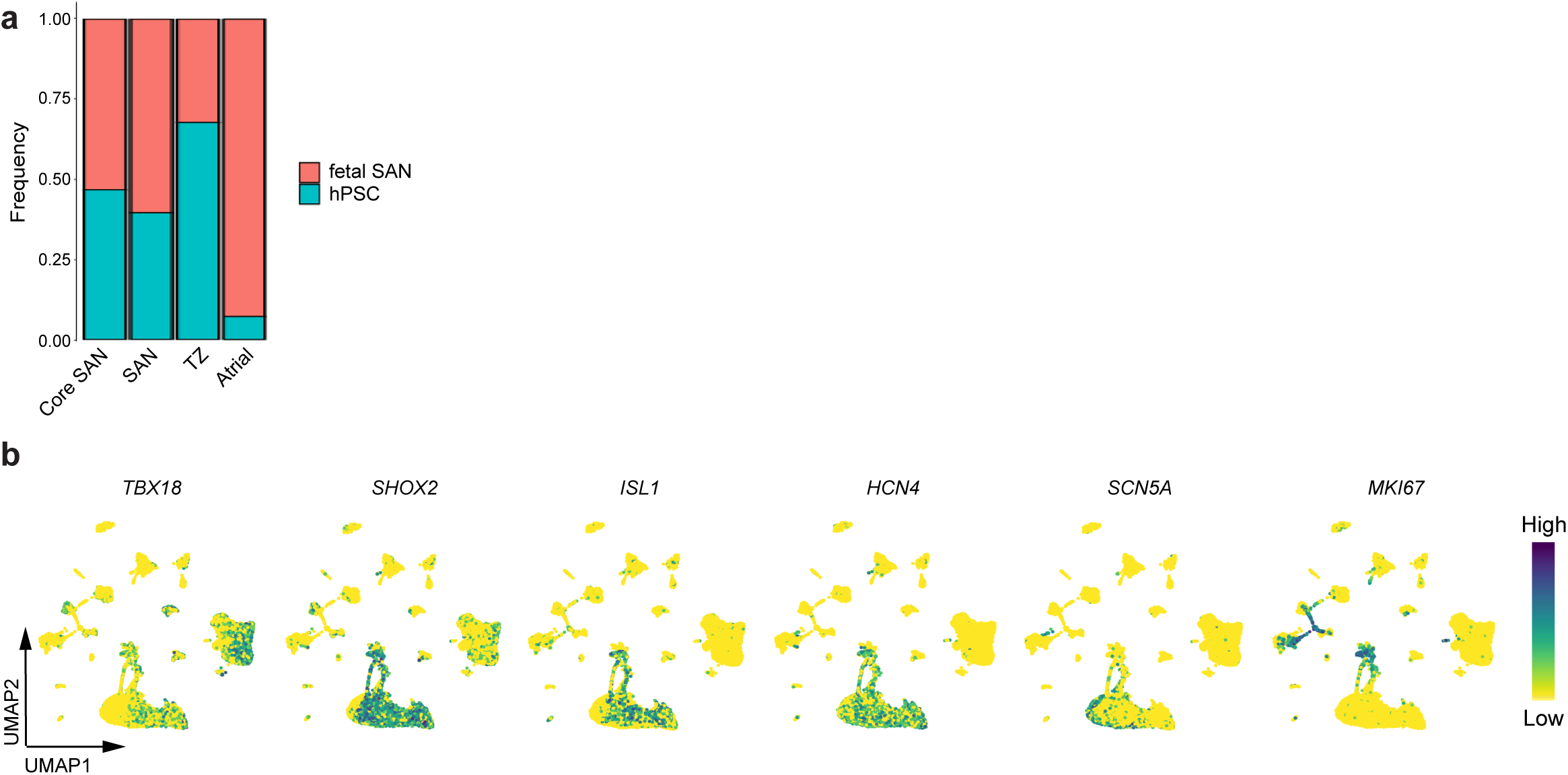
Single-cell RNA sequencing of hPSC-derived SANLPC reveals transcriptomic similarities to fetal SAN pacemaker cells. **a** Stacked bar graph showing the frequency of fetal and hPSC-derived cells in the indicated clusters. **b** UMAPs of integrated fetal SAN and hPSC datasets showing the expression of the indicated genes.

**Supplementary Figure 2 (related to Figure 3):**
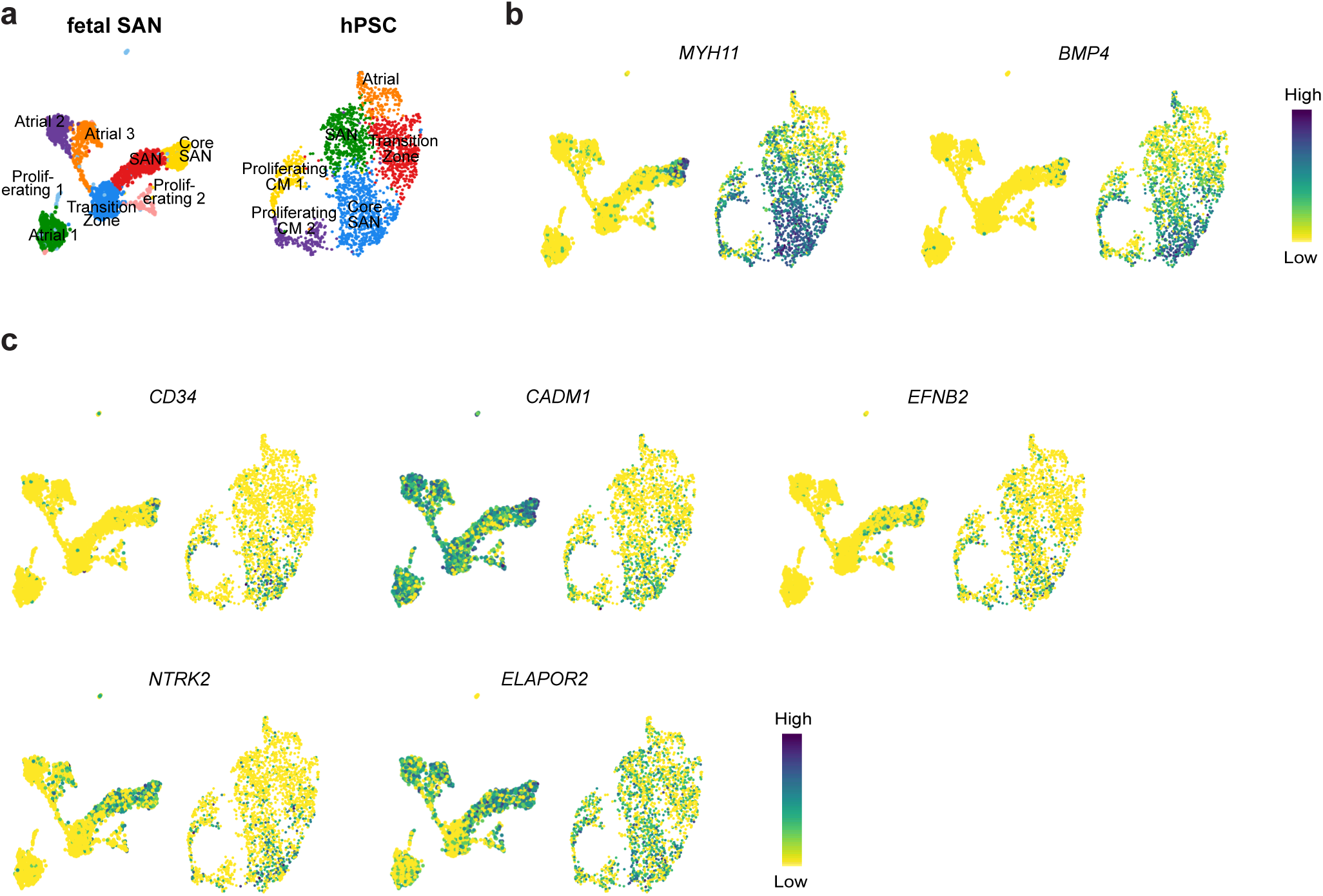
Comparison of fetal and hPSC-derived SAN cells identifies shared Core SAN markers. **a** UMAPs of fetal (left) and hPSC-derived (right) *TNNT2*^+^ cardiomyocytes showing the assigned cell types. **b** UMAPs of fetal (left) and hPSC-derived (right) *TNNT2^+^* cardiomyocytes showing the expression of the indicated genes. **c** UMAPs of fetal (left) and hPSC-derived (right) *TNNT2^+^* cardiomyocytes showing the expression of the five genes encoding for membrane spanning proteins that were identified in the shared Core SAN marker list.

**Supplementary Figure 3 (related to Figure 3):**
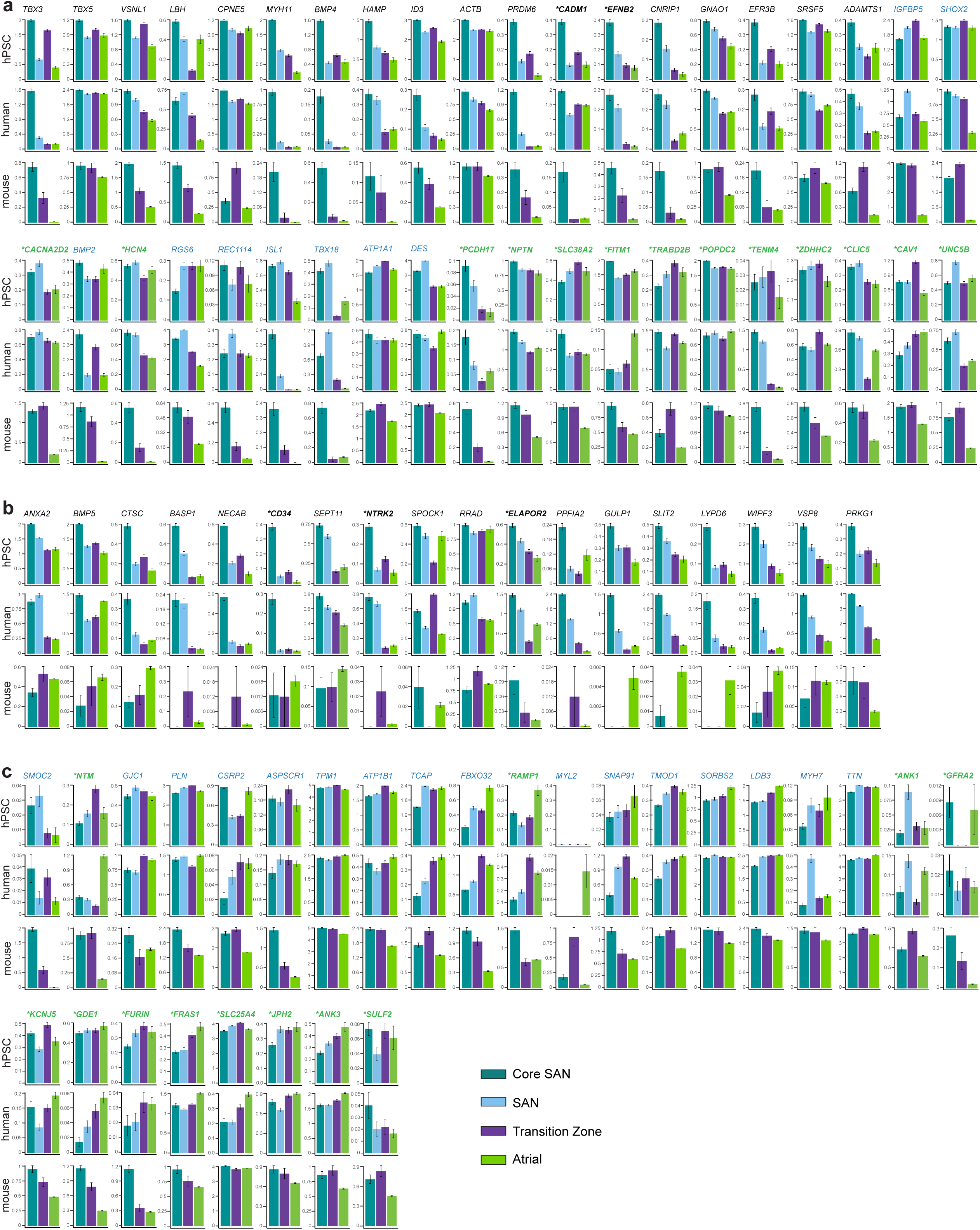
Identification of conserved and species-specific SAN pacemaker cell markers. **a** Bar graphs showing the expression of the indicated genes that are conserved markers of both human and mouse SAN pacemaker cells. **b** Bar graphs showing the expression of the indicated genes that are specifically expressed in human but not mouse SAN pacemaker cells. **c** Bar graphs showing the expression of the indicated genes that are specifically expressed in mouse but not human SAN pacemaker cells. hPSC, expression in hPSC- derived scRNA-seq dataset; human, expression in human fetal SAN snRNA-seq dataset; mouse, expression in mouse SAN scRNA-seq dataset from Goodyer et al.^9, 25^; genes labelled in black are contained in the list of 36 Core SAN marker genes identified in this study; genes labelled in blue were identified in the mouse by Goodyer et. al 2019^9^; genes labelled in green were identified as SAN surface markers in the mouse by Goodyer et. al 2022^25^; *indicates surface markers.

**Supplementary Figure 4 (related to Figure 3):**
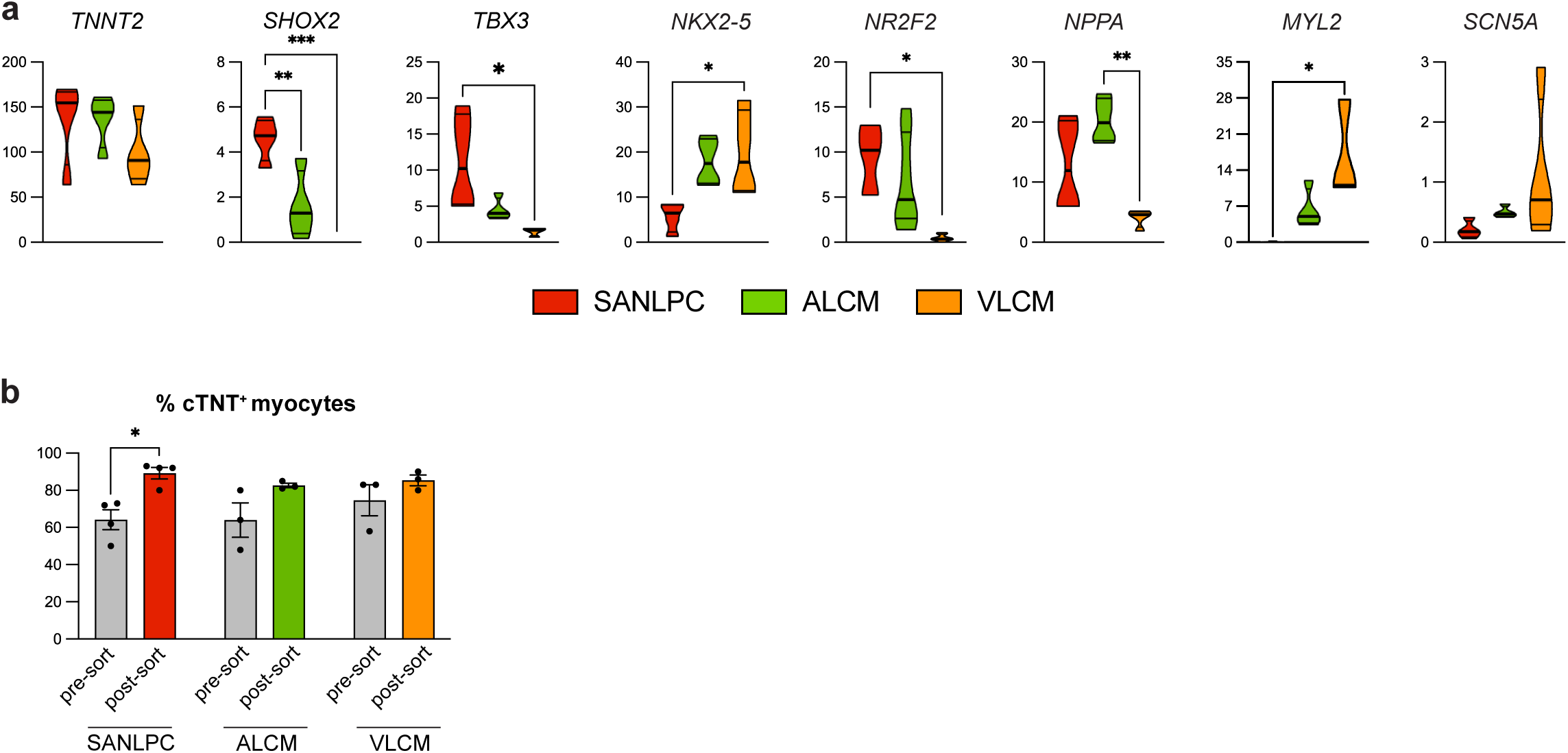
MYH11, BMP4 and CD34 are specifically expressed in hPSC-derived SANLPCs. **a** RT-qPCR analysis of the expression of the indicated genes in hPSC-derived SANLPCs, ALCMs and VLCMs at day 25 (n = 4). Values represent expression relative to the housekeeping gene TBP. **b** Bar graph showing the % of cTNT^+^ cells before and after enrichment for cardiomyocytes using the PSC-Derived Cardiomyocyte Isolation Kit from Miltenyi Biotec. The cardiomyocyte enriched cultures of SANLPCs, ALCMs and VLCMs were used for the immunofluorescent stainings shown in Figure 3e-h (n = 3). Statistical analysis was performed using one-way ANOVA followed by Bonferroni’s post hoc test: *p < 0.05, **p < 0.01, ***p < 0.001 vs indicated sample. Error bars represent SEM.

**Supplementary Figure 5 (related to Figure 4):**
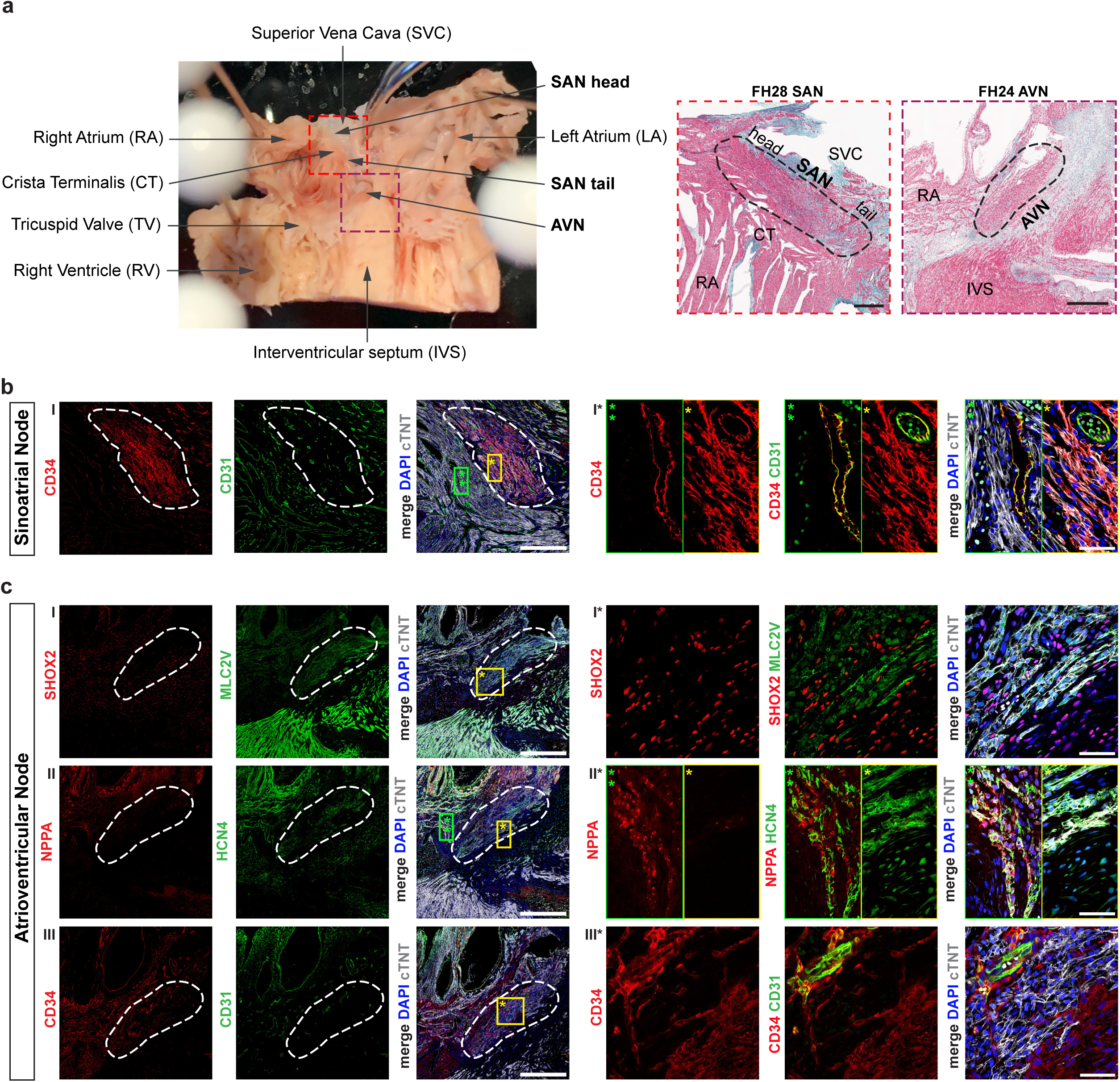
MYH11, BMP4 and CD34 are specifically expressed in human SAN pacemaker cardiomyocytes. **a** Image showing the dissection of a human fetal heart for the preparation of SAN and AVN tissue sections (left) and low magnification Masson’s trichrome staining of fetal heart #28 SAN (gestation week 19) and fetal heart #24 AVN (gestation week 17) tissue sections shown here and in Fig. 4 (right). Black dashed line outlines the SAN / AVN respectively. Scale bars, 500μm. **b** Immunofluorescent staining of gestation week 19 fetal human SAN for: CD34 and endothelial cell marker CD31 (I). White dashed line outlines the SAN. Yellow and green boxes indicate location of high magnification insets shown on the right marked with *. **c** Immunofluorescent staining of gestation week 17 fetal human AVN for: SAN pacemaker transcription factor SHOX2 and ventricular myocyte marker MLC2V (I), atrial myocyte marker NPPA and pacemaker ion channel HCN4 (II), and CD34 and endothelial cell marker CD31 (III). Images represent consecutive sections of AVN tissue. White dashed line outlines the AVN. Yellow and green boxes indicate location of high magnification insets shown on the right marked with *. Scale bars, 500μm (left) and 50μm in the high magnification insets (right).

**Supplementary Figure 6 (related to Figure 4):**
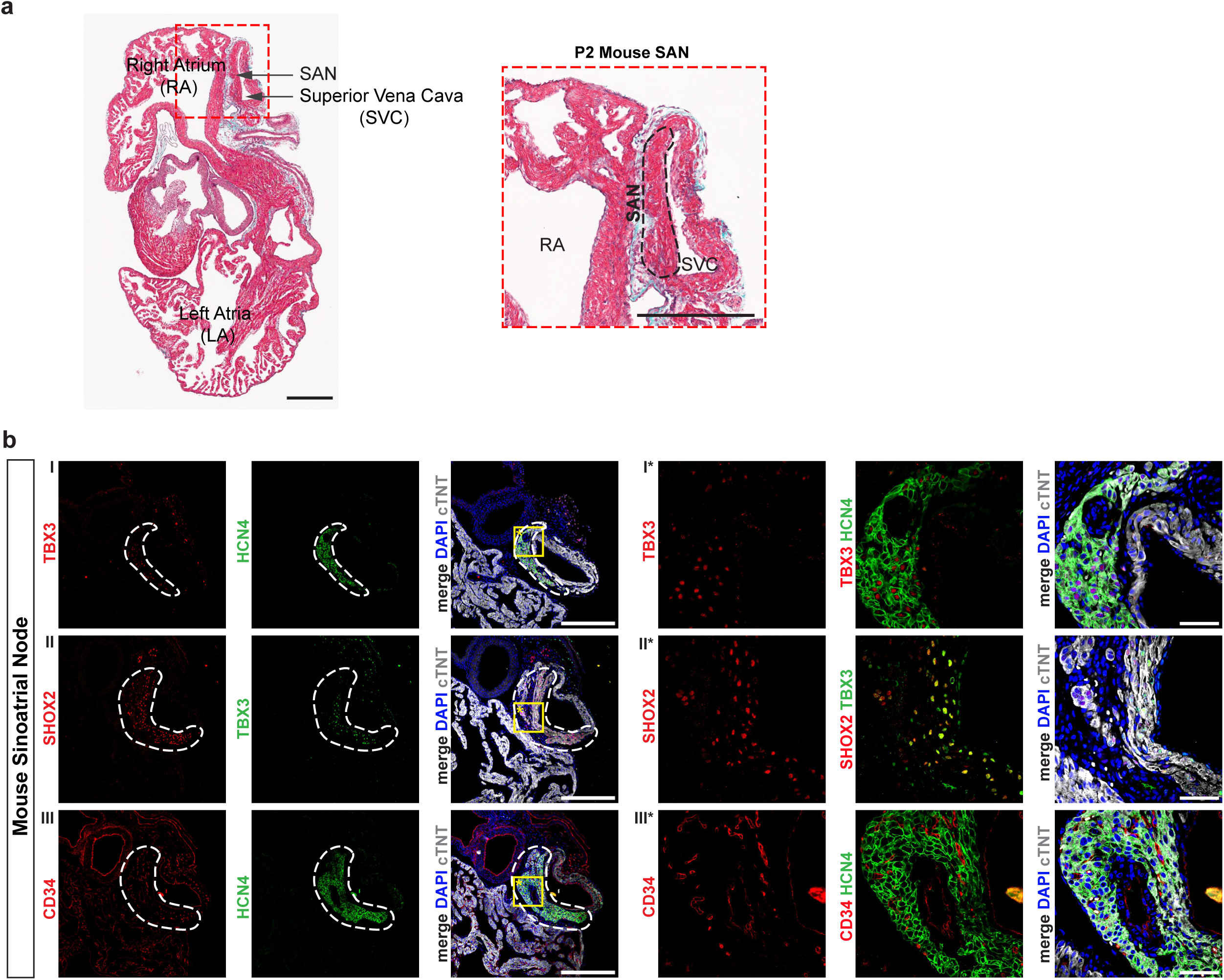
CD34 is not expressed in mouse SAN pacemaker cardiomyocytes. **a** Low magnification Masson’s trichrome staining of mouse postnatal day 2 heart section containing cross section of SAN tissue (left) and higher magnification inset of SAN tissue (right). Black dashed line outlines the SAN. Scale bars, 300μm. **b** Immunofluorescent staining of mouse SAN for: pacemaker transcription factor TBX3 and pacemaker ion channel HCN4 (I), SAN pacemaker transcription factors SHOX2 and TBX3 (II), and CD34 and HCN4 (III). Images represent consecutive sections of mouse hearts. White dashed line outlines the SAN. Yellow box indicates location of high magnification insets shown on the right marked with *. Scale bars, 300μm (left) and 50μm in the high magnification insets (right).

**Supplementary Figure 7 (related to Figure 5):**
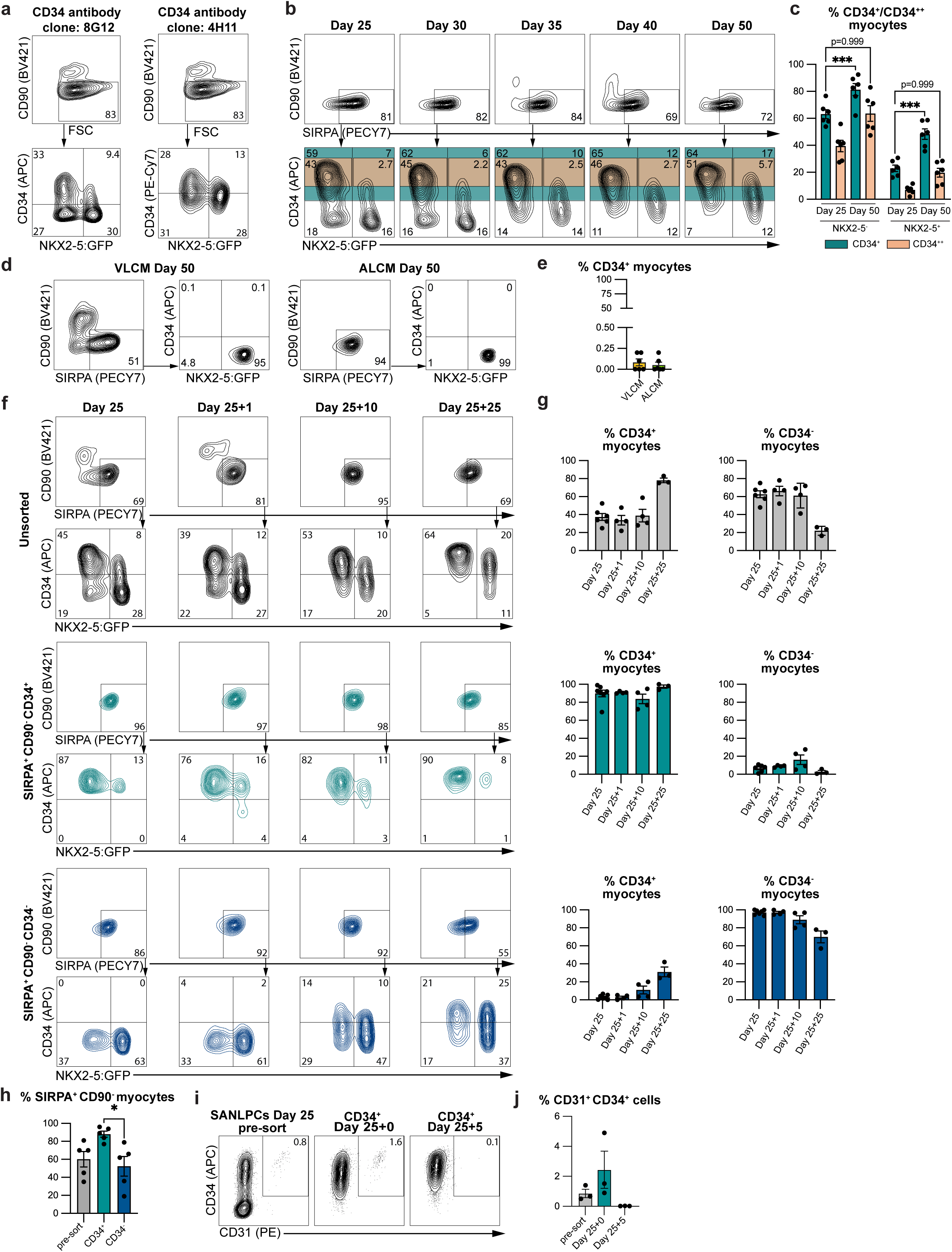
FACS and MACS sorting for CD34^+^ cells enriches for SANLPCs. **a** Flow cytometric analysis at day 25 of CD34 and NKX2-5:GFP expression in SIRPA^+^CD90^-^ cardiomyocytes in SANLPCs. Two different CD34 antibody clones were used to confirm consistent results for CD34 expression. **b** Flow cytometric analyses of CD34 and NKX2-5:GFP expression in SIRPA^+^CD90^-^ cardiomyocytes at indicated time points throughout the differentiation. Teal-colored gates indicate CD34^+^ cells and orange-colored gates indicate CD34^++^ cells. **c** Bar graph summarizing the expression of CD34 within the NKX2-5^-^ and NKX2-5^+^ myocyte fractions at day 25 and day 50 as shown in (b) (n = 6). **d** Flow cytometric analyses of CD34 and NKX2-5:GFP expression in SIRPA^+^CD90^-^ cardiomyocytes in VLCMs and ALCMs at day 50. **e** Bar graph summarizing the expression of CD34 in VLCMs and ALCMs as shown in (d) (n = 6). **f** Flow cytometric analyses of CD34 and NKX2-5:GFP expression in SIRPA^+^CD90^-^ cardiomyocytes in unsorted (top), CD34^+^ (centre), and CD34^-^ (bottom) FACS sorted samples at the indicated timepoints. **g** Bar graphs summarizing the proportion of CD34^+^ and CD34^-^ myocytes in unsorted (top), CD34^+^ (centre), and CD34^-^ (bottom) FACS sorted samples at the indicated timepoints as shown in (f) (n = 3-6). **h** Bar graph summarizing the proportion of SIRPA^+^CD90^-^ cardiomyocytes in the indicated populations before and after MACS of SANLPC cultures at day 25 (n = 5). **i** Flow cytometric analyses of CD31 and CD34 expression before and after MACS of SANLPC cultures at the indicated time points. **j** Bar graph summarizing the proportion of CD31^+^CD34^+^ endothelial cells at the indicated time points as shown in (i) (n = 3). Statistical analysis was performed using one-way ANOVA followed by Bonferroni’s post hoc test: *p < 0.05, **p < 0.01, ***p < 0.001 vs indicated sample. Error bars represent SEM.

**Supplementary Figure 8 (related to Figure 5):**
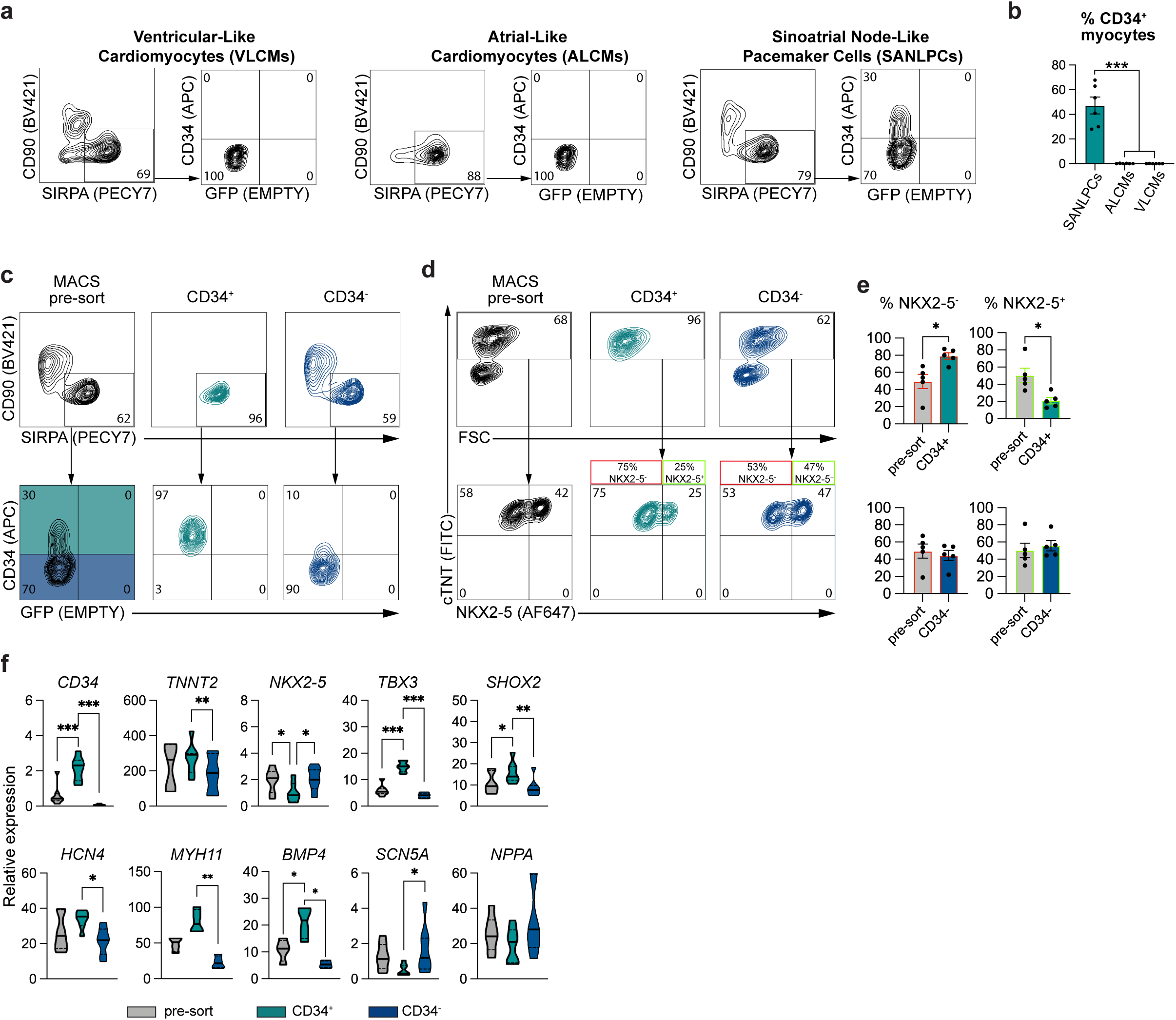
MACS sorting for CD34^+^ cells enriches for SANLPCs from the HES2 hPSC line. **a** Flow cytometric analyses at day 25 of CD34 expression in SIRPA^+^CD90^-^ cardiomyocytes in HES2-derived VLCMs, ALCMs, and SANLPCs. **b** Bar graphs summarizing the expression of CD34 in myocytes as shown in (a) in the indicated differentiation cultures. **c** Flow cytometric analyses of CD34 expression in SIRPA^+^CD90^-^ cardiomyocytes before and after MACS at day 25. Teal shading indicates CD34^+^ and blue shading indicates CD34^-^ sorting gates. **d** Flow cytometric analyses of NKX2-5 expression in cTNT^+^ cardiomyocytes before and after MACS. Note, because the HES2 cell line does not carry a NKX2-5:GFP reporter intracellular staining for NKX2-5 was performed. **e** Bar graphs summarizing the proportion of NKX2-5^-^ and NKX2-5^+^ cells in presort, CD34^+^, and CD34^-^ MACS sorted samples (n = 5). **f** RT-qPCR analysis of the expression of the indicated genes in presort, CD34^+^, and CD34^-^ MACS sorted samples (n = 4-7). Values represent expression relative to the housekeeping gene TBP. Statistical analysis was performed using two-sided unpaired t-test when comparing two samples (e) and one-way ANOVA followed by Bonferroni’s post hoc test when comparing >2 samples (b, f): *p < 0.05, **p < 0.01, ***p < 0.001 vs indicated sample. Error bars represent SEM.

**Supplementary Figure 9 (related to Figure 6):**
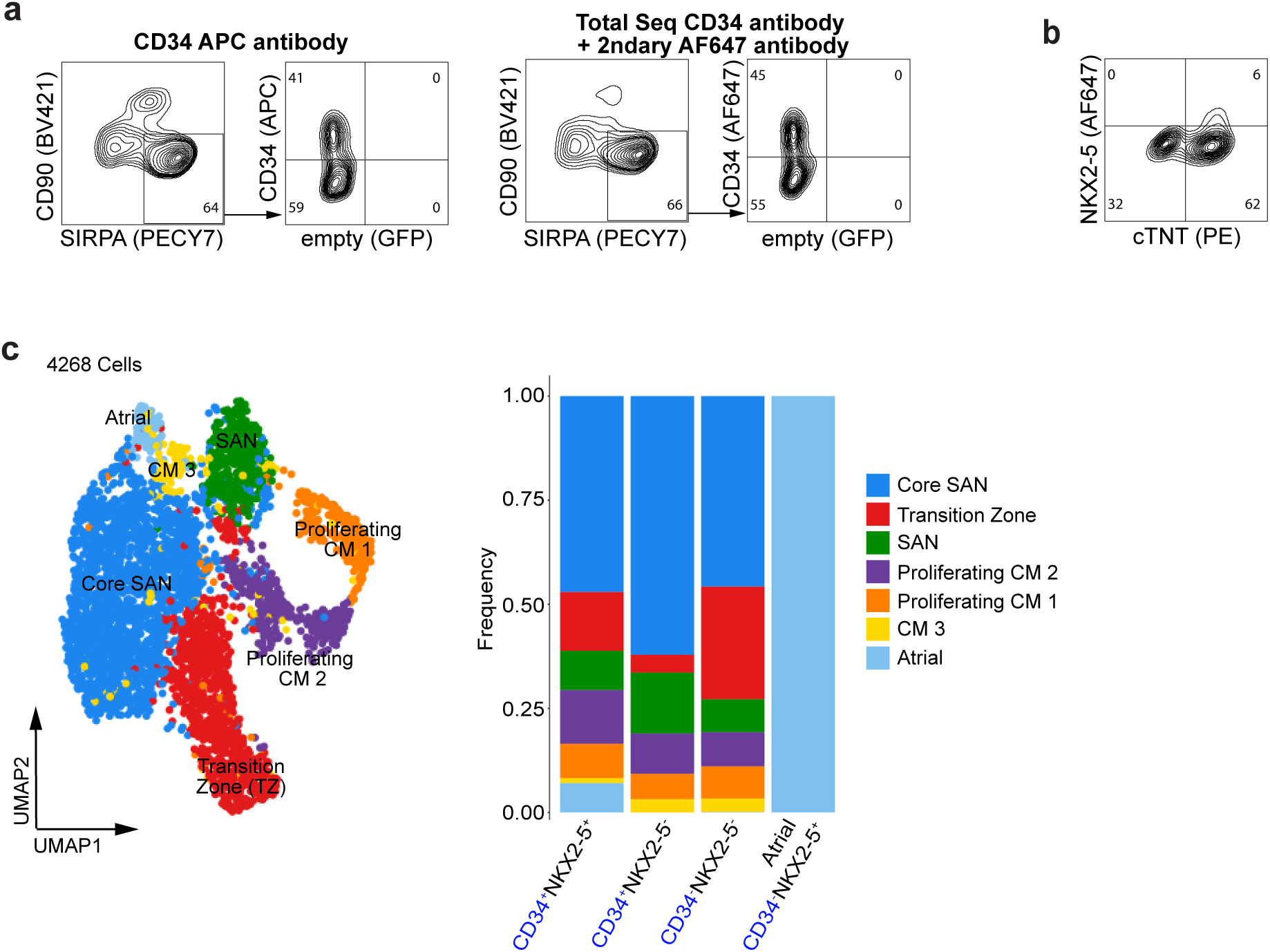
CITE-seq of hPSC-derived SANLPCs shows that CD34^+^NKX2-5^+^ and CD34^-^NKX2-5^-^ cardiomyocytes are transcriptionally similar to SAN pacemaker cells. **a** Flow cytometric analyses of the day 25 HES2 SANLPC differentiation cultures used for the CITE-Seq experiment showing expression of CD34 in SIRPA^+^CD90^-^ cardiomyocytes. Samples were stained with either the regular CD34-APC antibody (left) or the barcoded Total Seq CD34 antibody used to detect CD34 protein expression in the CITE-seq experiment (right). A secondary AF647 antibody against the Total Seq CD34 primary antibody was used to visualize the cells detected with the Total Seq antibody. Note, the proportion of CD34^+^ cells was comparable for both stains. **b** Flow cytometric analyses of the day 25 HES2 SANLPC differentiation cultures used for the CITE-Seq experiment showing the expression of cTNT and NKX2-5. **c** UMAP of the subclustered cardiomyocytes from the day 25 HES2-derived SANLPC CITE-seq dataset showing the assigned cell types (left). Stacked bar graph showing the frequency of each cell type in the indicated cell populations (right). Blue label indicates that protein level CD34 expression was used for the selection of the cell populations.

**Supplementary Figure 10:**
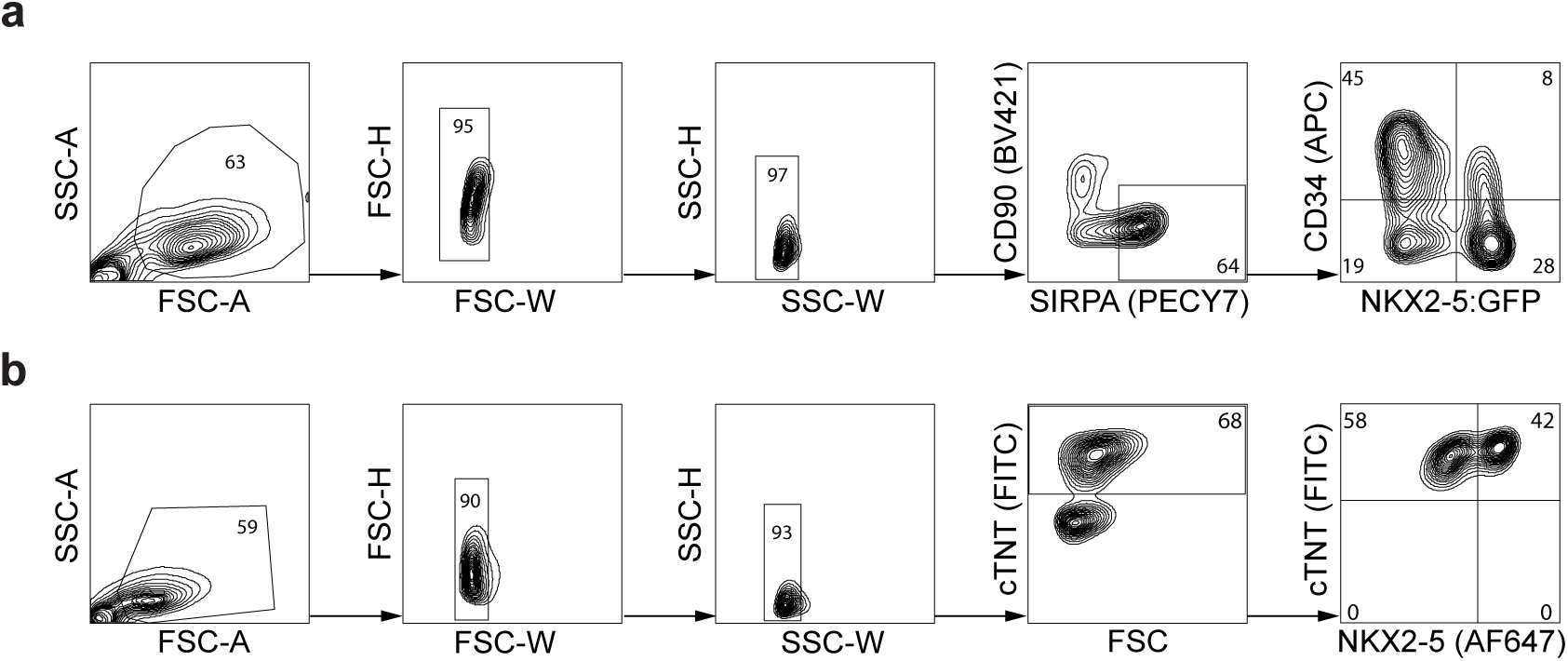
Gating strategies used for flow cytometric analyses. **a** Gating strategy used to determine the expression of CD34 and NKX2-5:GFP in SIRPA^+^CD90^-^ cardiomyocytes shown in Figure 5, and Supplementary Figures 7, 8, 9. **b** Gating strategy used to determine the expression of cTNT and NKX2-5 shown in Supplementary Figures 4, 8, 9.

